# Tracing the development and lifespan change of population-level structural asymmetry in the cerebral cortex

**DOI:** 10.1101/2021.11.25.469988

**Authors:** James M. Roe, Didac Vidal-Piñeiro, Inge K. Amlien, Mengyu Pan, Markus H. Sneve, Michel Thiebaut de Schotten, Patrick Friedrich, Zhiqiang Sha, Clyde Francks, Yunpeng Wang, Kristine B. Walhovd, Anders M. Fjell, René Westerhausen

## Abstract

Cortical asymmetry is a ubiquitous feature of brain organization that is subtly altered in some neurodevelopmental disorders, yet we lack knowledge of how its development proceeds across life in health. Achieving consensus on the precise cortical asymmetries in humans is necessary to uncover the genetic and later influences that shape them, such as age. Here, we delineate population-level asymmetry in cortical thickness and surface area vertex-wise in 7 datasets and chart asymmetry trajectories longitudinally across life (4-89 years; observations = 3937; 70% longitudinal). We find replicable asymmetry interrelationships, heritability maps, and test asymmetry associations in large-scale data. Cortical asymmetry was robust across datasets. Whereas areal asymmetry is predominantly stable across life, thickness asymmetry grows in childhood and peaks in early adulthood. Areal asymmetry correlates phenotypically and genetically in specific regions, and is low-moderately heritable (max h^2^_SNP_ ∼19%). In contrast, thickness asymmetry is globally interrelated across the cortex in a pattern suggesting highly left-lateralized individuals tend towards left-lateralization also in population-level right-asymmetric regions (and vice versa), and exhibits low or absent heritability. We find less areal asymmetry in the most consistently lateralized region in humans associates with subtly lower cognitive ability, and confirm small handedness and sex effects. Results suggest areal asymmetry is developmentally stable and arises in early life through genetic but mainly subject-specific stochastic effects, whereas childhood developmental growth shapes thickness asymmetry and may lead to directional variability of global thickness lateralization in the population.

## 1. Introduction

The brain’s hemispheres exhibit high contralateral symmetry ^1,2^, homotopic regions are under similar genetic influence ^3–5^ and show highly correlated developmental change ^3,6^. Despite this, structural asymmetry is also a ubiquitous aspect of brain organization ^7,8^. Cortical thickness and surface area are known to exhibit distinct asymmetry patterns ^7,9^, albeit reported inconsistently ^7,8,18–22,10–17^. Yet achieving consensus on cortical asymmetries in humans is a prerequisite to uncover the genetic-developmental and lifespan influences that shape and alter them. Although an extensive literature in search of structural asymmetry deviations in various conditions and disorders is in several cases being challenged by newer data ^23^, at least some aspects of cortical asymmetry are confirmed to be subtly reduced in neurodevelopmental disorders such as autism ^24,25^, but also through later life influences such as aging ^18^, and Alzheimer’s disease ^18,26^. Hence, altered cortical asymmetry at various lifespan stages may be associated with reduced brain health. However, it is currently unknown how cortical asymmetry development proceeds across life in health, because no previous study has charted cortical asymmetry trajectories from childhood to old age using longitudinal data.

Compounding the lack of longitudinal investigation, previous large-scale studies do not delineate the precise brain regions exhibiting robust cortical asymmetry ^7^, relying on brain atlases with predefined anatomical boundaries that may not conform well to the underlying asymmetry of cortex ^7,27^. Taking an atlas-free approach to delineate asymmetries that reliably reproduce across international samples as starting point (i.e. population-level asymmetries) would better enable mapping of the developmental principles underlying structural cortical asymmetries, as well as the genetic and individual-specific factors associated with cortical lateralization. Furthermore, such an approach would help resolve the many reported inconsistencies for cortical asymmetry maps ^7,8,18–21,10–17^ – e.g. reports of both right- and left-thickness lateralization in medial (^11,13,14,28^ and ^17–20^) and lateral prefrontal cortex (PFC; ^17,18,20,22,28^ and ^8,10–12,16^), and right-^11,31^ and left-^7,29,30^ areal lateralization of superior temporal sulcus (STS) – while serving as a high-fidelity phenotype for future brain asymmetry studies to complement existing low-resolution atlases ^7^.

Determining the developmental and lifespan trajectories of cortical asymmetry may shed light on how cortical asymmetries are shaped through childhood or set from early life, and provide evidence of the timing of expected brain change in normal development. Although important in and of itself, this would also provide a useful normative reference, as subtly altered cortical asymmetry – in terms of both area and thickness – has been linked at the group-level to neurodevelopmental disorders along the autism spectrum ^24,25^, suggesting altered lateralized neurodevelopment may be a neurobiologically relevant outcome in at least some cases of developmental perturbation ^24,25^. For areal asymmetry, surprisingly few studies have charted developmental ^30,31^ or aging-related effects ^7,28^, although indirect evidence in neonates suggests adult-like patterns of areal asymmetry are evident at birth and may exhibit little change from birth to 2 years ^31^ despite rapid and concurrent developmental cortical expansion ^32^. For thickness asymmetry, longitudinal increases in asymmetry have been shown during the first two years of life ^19^, with suggestions of rapid asymmetry growth from birth to 1 year ^19^, and potentially continued growth until adolescence ^33^. However, previous lifespan studies mapped thickness asymmetry linearly across cross-sectional developmental and adult age-ranges ^10,17^, mostly concluding thickness asymmetry is minimal in infancy and maximal age ∼60. In contrast, recent work established thickness asymmetry shows a non-linear decline from 20 to 90 years that is reproducible across longitudinal aging cohorts ^18^. Thus, although offering viable developmental insights ^10,17^, previous lifespan studies of thickness asymmetry do not accurately capture the aging process, and likely conflate non-linear developmental and aging trajectories with linear models. A longitudinal exploration of the lifespan trajectories of thickness asymmetry accounting for dynamic change is needed to further knowledge of normal human brain development.

Correlations between cortical asymmetries may provide a window on asymmetries formed under common genetic or developmental influences. Contemporary research suggests brain asymmetries are complex, multifactorial and largely independent (i.e. uncorrelated) traits ^34–36^, contrasting earlier theories emphasizing a single ^37,38^ or predominating factor ^39,40^ controlling various cerebral lateralizations. Yet while there has been much research on whether asymmetries of various morphometric measures ^8,11,16^ or imaging modalities ^29^ relate to one another, few have focused on interregional relationships between asymmetries derived from the same metric. Where reported, evidence suggests cortical asymmetries may be mostly independent ^27,41,42^ – in line with a multifactorial view ^34,35,43^ – and a recent study found asymmetry in anatomically connected regions of the cortical language network was no more related than in regions selected at random ^29^. Currently, it is not known whether or how cortical asymmetries may correlate within individuals, though this may signify coordinated genetic or later development of left-right brain asymmetries – a genetic or later developmental account depending on joint evidence of trait heritability, and whether or not phenotypic correlations are underpinned by genetic correlations.

Finally, altered development of cerebral lateralization in general has been widely hypothesized to relate to average poorer cognitive outcomes ^17,44,45^. Specifically in the context of cortical asymmetry, however, although one previous study reported larger thickness asymmetry may relate to better verbal and visuospatial cognition ^17^, phenotypic asymmetry-cognition associations have been rarely reported ^17,46,47^, conflicting ^46,47^, not directly comparable ^17,46^, and to date remain untested in large-scale data. Still, recent work points to small but significant overlap between genes underlying multivariate brain asymmetries and those influencing educational attainment and specific developmental disorders impacting cognition ^27^, indicating either pleiotropy between non-related traits or capturing shared genetic susceptibility to altered brain lateralization and cognitive outcomes. Furthermore, most large-scale studies of the factors widely assumed to be important in the context of asymmetry have adopted brain atlases offering limited spatial precision ^7,23,27^. Accordingly, such studies did not detect associations with handedness ^7,48^ that were not found until a recent study applied higher resolution (i.e. vertex-wise) mapping in big data ^22^. Therefore, as a final step, we reasoned that combining an optimal delineation of population-level cortical asymmetries with big data would enhance detection and quantification of such effects – i.e. general cognitive ability, handedness and sex.

Here, we first delineate population-level cortical areal and thickness asymmetries using vertex-wise analyses and their overlap in 7 international datasets. To gain insight into their development, we then trace a series of lifespan and genetic analyses. Specifically, we chart the developmental and lifespan trajectories of cortical asymmetry for the first time longitudinally across the lifespan. We then examine phenotypic interregional asymmetry correlations – under the assumption correlations indicate coordinated development of left-right asymmetries through genes or lifespan influences – test heritability using both extended twin and genome-wide single nucleotide polymorphism (SNP) data, and whether phenotypic associations are underpinned by genetics. Finally, we screened our set of robust, population-level asymmetries for association with general cognitive ability and factors purportedly related to asymmetry in UK Biobank (UKB) ^49^. Based on findings of aging-related dedifferentiation in thickness asymmetry ^18^, we hypothesized trajectories of cortical thickness would show developmental growth in thickness asymmetry (i.e. differentiation), but remained agnostic regarding lifespan areal asymmetry development.

## 2. Results

### 2.1 Population-level asymmetry of the cerebral cortex

First, to delineate cortical regions exhibiting population-level areal and thickness asymmetry, we assessed asymmetry vertex-wise in 7 independent adult samples and quantified overlapping effects (Methods). Areal asymmetries were highly consistent across all 7 datasets (Figure 1A): the spatial correlation between surface AI maps ranged from *r* = .88 to .97 (Figure 1C). Across all 7 datasets (Figure 1D), overlapping effects for strong leftward areal asymmetry were observed in a large cluster in supramarginal gyrus (SMG) that spanned postcentral gyrus, planum temporale and primary auditory regions (and conformed well to their anatomical boundaries; see Figure 1–figure supplement 1A for significance), anterior insula and temporal cortex, rostral anterior cingulate, medial superior frontal cortex, and precuneus, the latter spanning parahippocampal and entorhinal cortex. Overlapping effects for strong rightward areal asymmetry were evident in cingulate, inferior parietal cortex, STS, medial occipital cortex, mPFC and rostral middle frontal cortex (Figure 1D). This global pattern agrees with previous reports ^7,21,22^.

**Figure 1.**
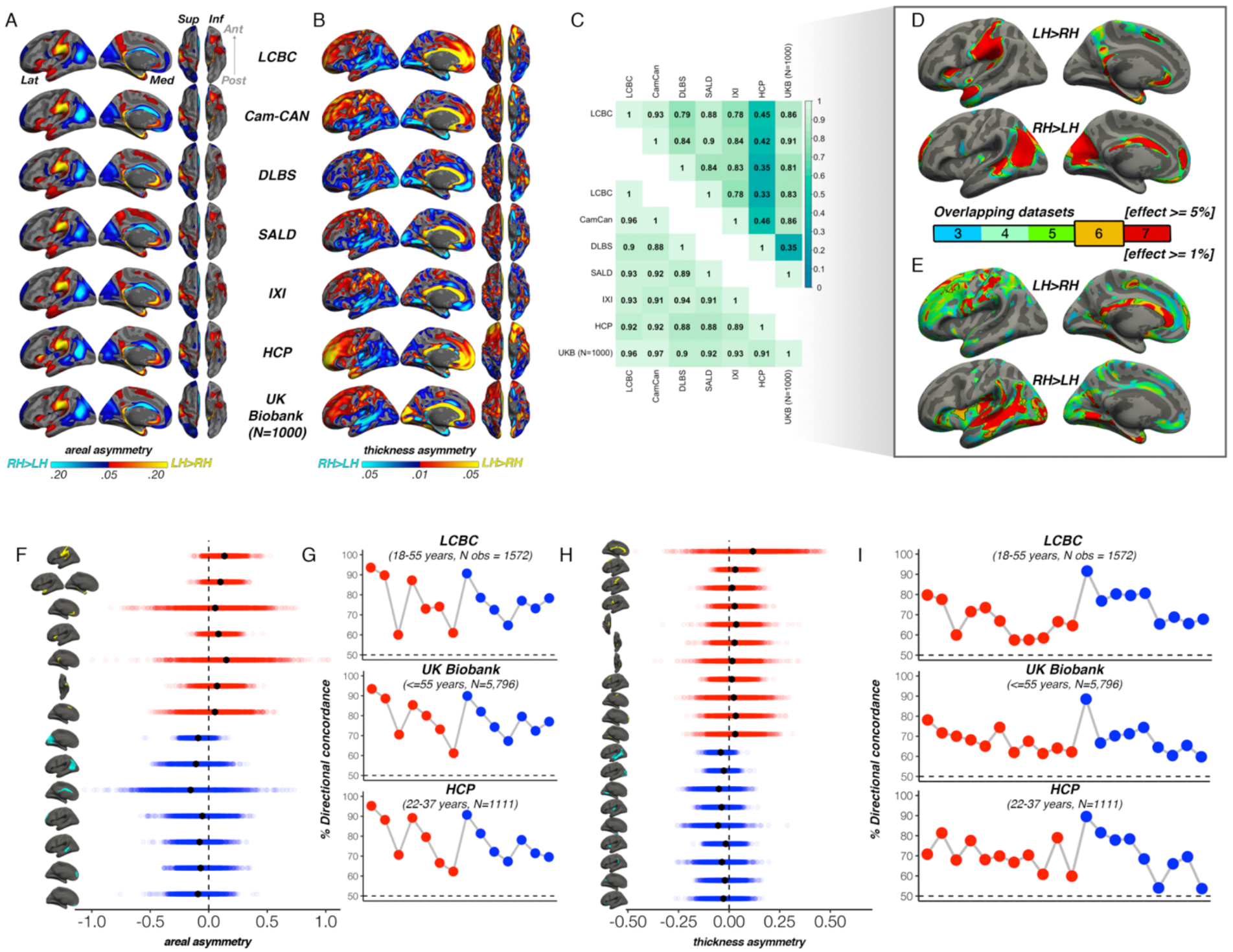
**A)** Mean areal and **B)** thickness asymmetry in each dataset. Warm and cold colours depict leftward and rightward asymmetry, respectively. **C)** Spatial overlap (Pearson’s r) of the unthresholded maps between datasets for areal (lower matrix) and thickness asymmetry (upper). **D)** Overlapping effects across datasets were used to delineate clusters exhibiting population-level areal (lower threshold = 5%) and **E)** thickness asymmetry (lower threshold = 1%) based on a minimum 6-dataset overlap (black outlined clusters). **F, H)** Raw distribution of the individual-level asymmetry index (AI) in adults extracted from clusters exhibiting areal and thickness asymmetry, respectively. Mean AI’s are in black, Raw distributions are shown for the LCBC (18-55 years) dataset with mixed effects data (outliers defined in lifespan analysis removed on a region-wise basis; Methods; Supplementary File 1E-F). X-axis denotes the AI of the average thickness and area of a vertex within the cluster. **G, I)** Proportion of individuals with the expected directionality of asymmetry within each cluster exhibiting areal and thickness asymmetry, respectively, shown for the three largest adult datasets. The X-axes in G and D are ordered according to the clusters shown in F and H, respectively. Lat=lateral; Med=medial; Post=posterior; Ant=anterior; Sup=superior; Inf=inferior.

For thickness, an anterior-posterior pattern of left-right asymmetry was evident in most datasets (Figure 1B), consistent with more recent reports ^7,17,18,22,28^. Though spatial correlations between AI maps from independent datasets were high, they were notably more variable (*r* = .33 - .93; Figure 1C); HCP showed lower correlation with all datasets (*r* = .33 - .46) whereas all other datasets correlated highly (min *r* = .78). Consistent leftward thickness asymmetry effects were evident in cingulate, postcentral gyrus, and superior frontal cortex (Figure 1E), whereas consistent effects for rightward thickness asymmetry were evident in a large cluster in and around STS (Fig 1E; Figure 1–figure supplement 1B), insula, lingual gyrus, parahippocampal and entorhinal cortex. Of note, both areal and thickness asymmetry extended beyond these described overlapping effects (Figure 1–figure supplements 1-2 & 5).

Based on effect size criteria (Figure 1D-E; Methods), we derived a set of robust clusters exhibiting population-level areal (14 clusters) and thickness asymmetry (20 clusters) for further analyses (see Supplementary file 1E-F for anatomical descriptions). The proportion of individuals lateralized in the population direction in each cluster was highly similar across datasets, on average ranging between 61-94% for area, and 57-90% for thickness (Figure 1F-I). We then formally compared our approach to asymmetry estimates derived from a gyral-based atlas often used to assess asymmetry ^7,16,27^, finding fairly poor correspondence with the vertex-wise structure of cortical asymmetry for atlas-based regions, particularly for thickness asymmetry (Figure 1–figure supplement 3).

### 2.2 Lifespan trajectories of population-level cortical asymmetries

Having delineated regions exhibiting population-level areal and thickness asymmetry, we aimed to characterize the developmental and lifespan trajectories of cortical asymmetry from early childhood to old age, using a lifespan sample incorporating dense longitudinal data (Methods). For this, we used the mixed-effects LCBC lifespan sample covering the full age-range (4-89 years). To account for non-linear lifespan change, we used Generalized Additive Mixed Models (GAMMs) to model the smooth left- (LH) and right hemisphere (RH) age-trajectories in our robust clusters (Methods).

In all clusters, the homotopic areal trajectories revealed areal asymmetry was strongly established already by age ∼4, and the lifespan trajectories of both leftward (Figure 2A) and rightward (Figure 2B) asymmetries were largely parallel. Specifically, a large left-asymmetric region in and around SMG/perisylvian (#1; Figure 2A) showed strong asymmetry by age ∼4 that was largely maintained throughout life through steady aging-associated decline of both hemispheres, whereas leftward asymmetry of temporal cortex (#2,6) and anterior insular (#4) was maintained through developmental expansion and aging-associated decline of both hemispheres. Others (retrosplenial #5; mPFC #3,7) showed growth from pre-established asymmetry and more variable lifespan trajectories. On the other side, rightward asymmetries showed largely preserved asymmetry through aging-associated decline of both hemispheres (Figure 2B; medial occipital #1; lateral parietal #2; STS #5; orbitofrontal #7), through bilateral developmental expansion and aging-associated decline (mPFC #6), or steadily expanding bilateral surface area until mid-life (cingulate; #3). There was also little indication of relative hemispheric differences during cortical developmental expansion from 4-30 years (Figure 3–figure supplement 3A) or aging from 30-89 years (Figure 3–figure supplement 5). Though lifespan areal asymmetry trajectories did show significant change at some point throughout life in most clusters (Supplementary file 1E), factor-smooth GAMM interaction analyses confirmed that areal asymmetry was significantly different from zero across the entire lifespan in all clusters (Figure 3–figure supplements 1-2), and the average trajectories across all leftward and rightward clusters were clearly parallel (though still both exhibited a significant difference; bordered plots in Figure 2A-B; Supplementary file 1E).

**Figure 2:**
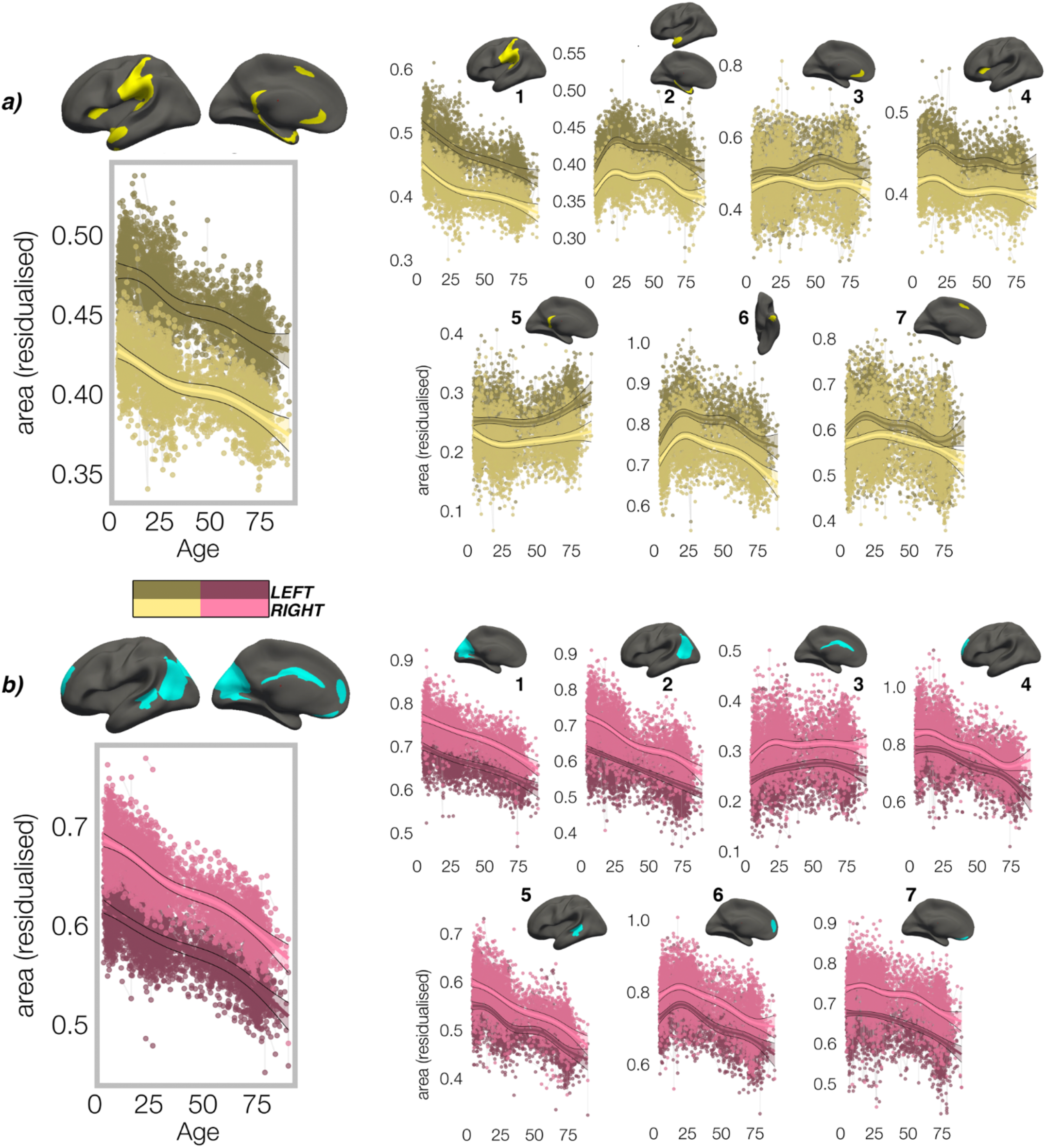
Homotopic lifespan trajectories of surface area in clusters exhibiting population-level **A)** leftward (yellow plots; yellow clusters) and **B)** rightward (pink plots; blue clusters) areal asymmetry (mm^2^). Larger plots on the left show the mean age trajectory across all clusters exhibiting leftward (top) and rightward (bottom) asymmetry. Note that the unit of measurement is the average surface area of a vertex within the cluster. Dark colours correspond to LH trajectories. All age trajectories were fitted using GAMMs. Data is residualized for sex, scanner and random subject intercepts. Clusters are numbered for reference.

In contrast, though homotopic trajectories of thickness clusters were more variable, they were non-parallel, and mostly characterized by developmental increase in thickness asymmetry from age 4-30, through seemingly unequal rates of continuous thinning between hemispheres (Figure 3; Figure 3–figure supplements 1-3). Importantly, in 10/20 clusters the data indicated developmental increase in thickness asymmetry corresponded to a significant relative hemispheric difference in the rate of developmental thinning. Mostly, these conformed to a pattern whereby the thicker homotopic hemisphere thinned comparatively slower (Figure 4): leftward thickness asymmetry developed through comparatively slower thinning of the LH that was significant in 6 clusters (superior, lateral and medial PFC #2 #8 #10, precentral #4, inferior temporal #6, calcarine #11; Figure 3A; Figure 4), whereas rightward asymmetry developed through significantly slower RH thinning (STS #1, planum temporale #7, anterior insula #9; Figure 3A; Figure 4), or significantly faster RH thickening (#5 entorhinal). Only one other cluster exhibited a relative hemispheric difference seemingly driven by faster thinning of the thicker hemisphere (#8 posterior cingulate; Figure 4). In these clusters, asymmetry development was generally evident until a peak in early adulthood (median age at peak = 24.3; see Figure 4) for both leftward and rightward clusters, around a point of inflection to less developmental thinning (see also Figure 3–figure supplement 4). The average trajectories across all leftward and rightward clusters also indicated developmental asymmetry increase (bordered plots; Figure 3). Despite the developmental growth, factor-smooth GAMMs nevertheless confirmed the developmental foundation for thickness asymmetry was already established by age ∼4 (95% of clusters exhibited small but significant asymmetry at age ∼4; Figure 3–figure supplement 2B), and again asymmetry trajectories showed significant change at some point throughout life (Supplementary file 1F). Across clusters delineated here we observed little evidence aging-related change from 30-89 years corresponded to a relative hemispheric difference in the rate of aging-related thinning, except in regions overlapping with our previous report ^18^ (e.g. medial PFC #10; Figure 3–figure supplement 5B; Figure 3–figure supplement 4). Thus, across population-level thickness asymmetries, the data indicated either developmental growth in asymmetry, or conserved relative asymmetry through development and aging despite absolute asymmetry change. Results were robust to varying the number of knots used to estimate trajectories (Figure 3–figure supplement 1).

**Figure 3:**
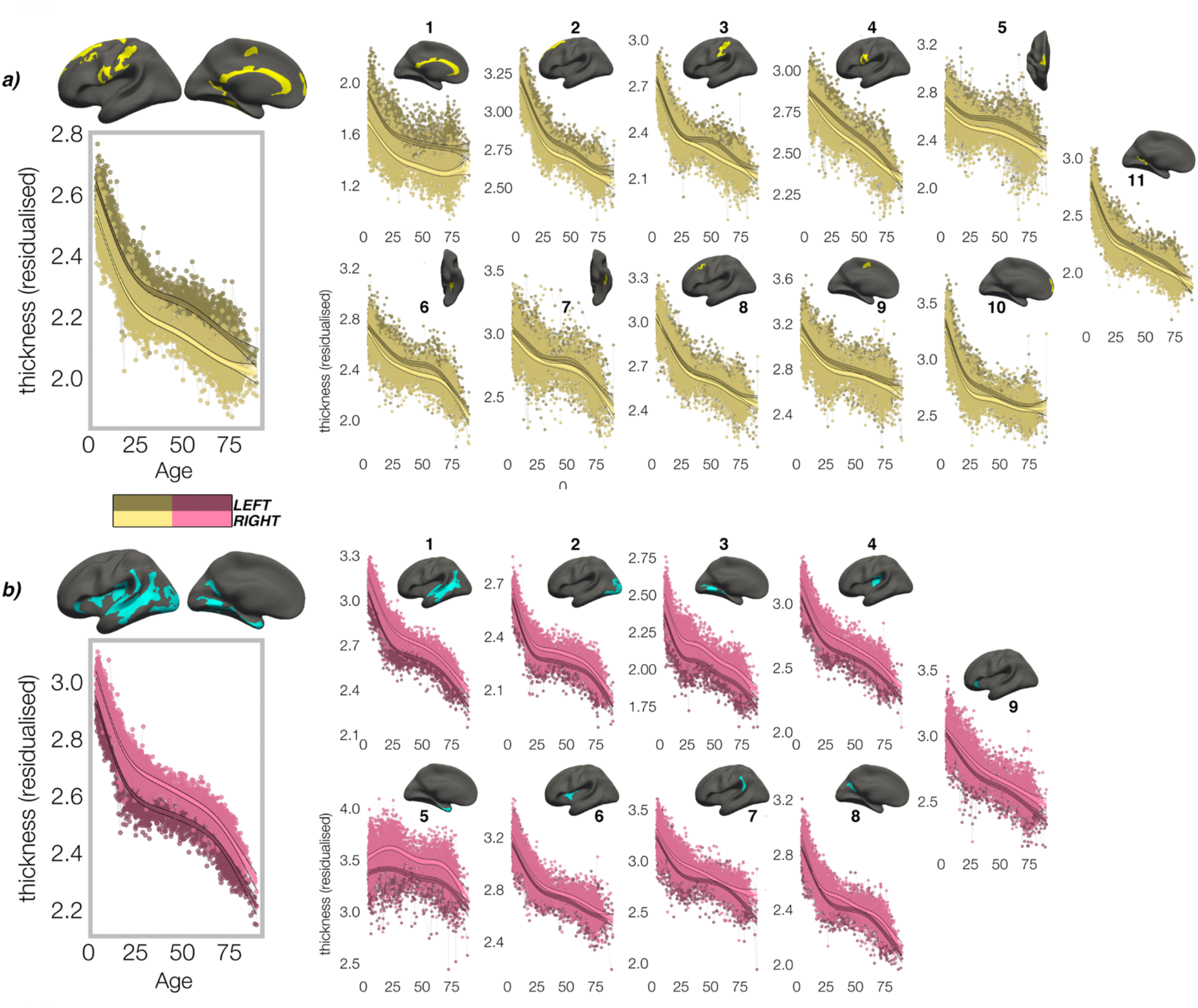
Homotopic lifespan trajectories of cortical thickness in clusters exhibiting population-level **A)** leftward (yellow plots; yellow clusters) and **B)** rightward (pink plots; blue clusters) thickness asymmetry (mm). Larger plots on the left show the mean age trajectory across all clusters exhibiting leftward (top) and rightward (bottom) asymmetry. Dark colours correspond to LH trajectories. All age trajectories were fitted using GAMMs. Data is residualized for sex, scanner and random subject intercepts. Clusters are numbered for reference.

**Figure 4:**
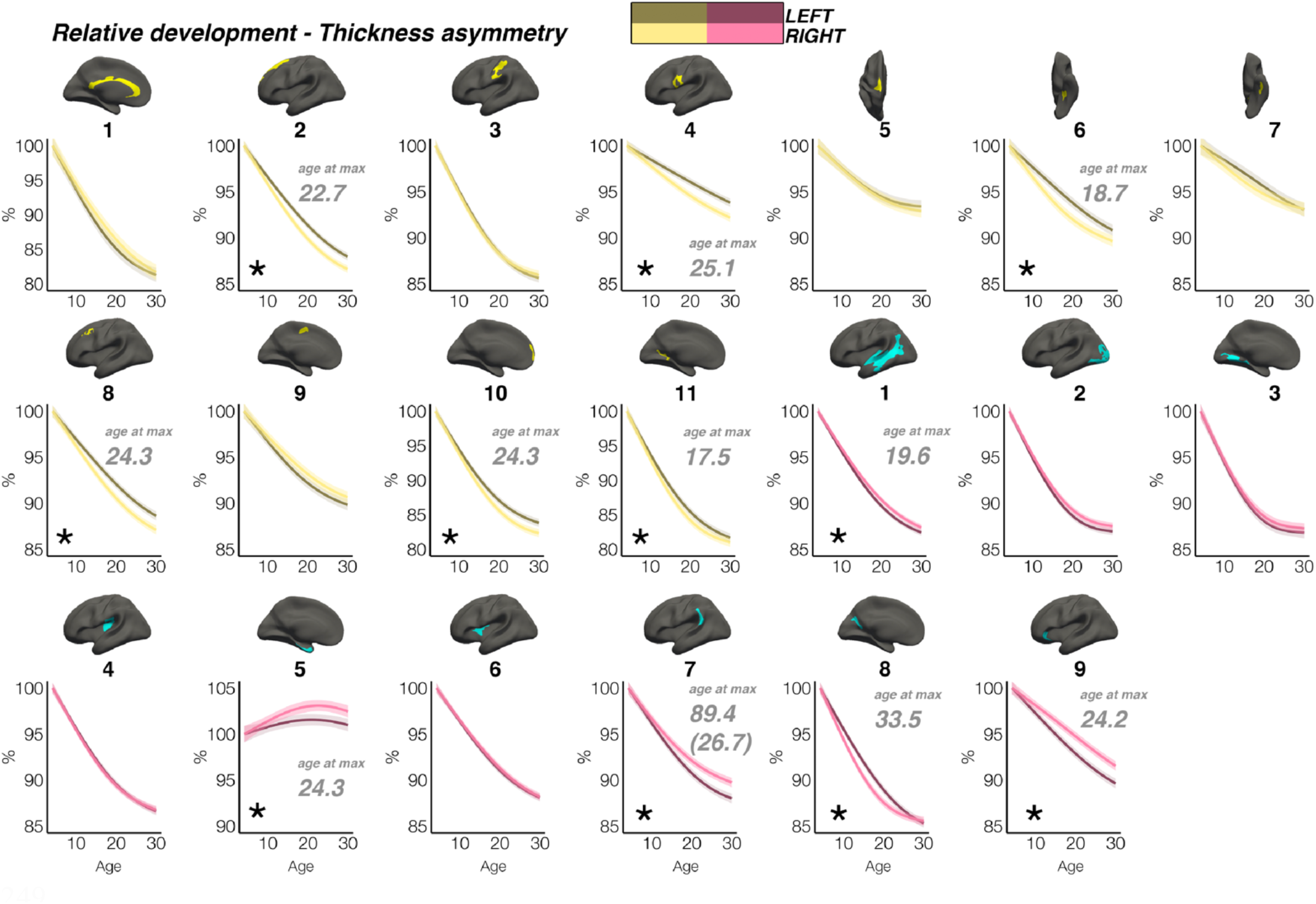
Relative developmental trajectories of homotopic cortical thickness in clusters exhibiting population-level thickness asymmetry. To highlight development, the x-axis covers the age-range 4-30 years, although relative change was calculated from full lifespan models (Methods). Darker trajectories indicate LH trajectories. Shaded areas indicate 95% CI. A relative difference in developmental thinning rates between hemispheres is suggested if the CI’s of relative LH and RH trajectories diverge to be non-overlapping (denoted with *). These indications should be interpreted in combination with the full lifespan GAMM trajectories shown in Figure 3. Where relative hemispheric differences are indicated, age at the point of maximum thickness asymmetry across life is denoted in grey (Methods). Since one cluster (R7) was estimated to exhibit maximum thickness asymmetry at age 89.4 (see Figure 3), age at maximum asymmetry for the developmental peak is also given in parentheses (Figure 3).

### 2.3 Interregional asymmetry correlations

We then investigated whether cortical asymmetry correlates within individuals (AI’s corrected for age, sex, scanner; Methods). For areal asymmetry, a common covariance structure between asymmetries was detectable across datasets: Mantel tests revealed the correlation matrices derived independently in LCBC, UKB and HCP data all correlated almost perfectly (*r* >= 0.97, all *p* < 9.9^−5;^ Figure 5A; Figure 5–figure supplement 1A). The highest correlations (or “hotspots”) all reflected positive correlations between regions that are on average left-asymmetric and regions that are on average right-asymmetric (i.e. higher leftward asymmetry in one region related to higher rightward asymmetry in another; Figure 5A black outline); leftward asymmetry in SMG/perisylvian (#1L) was related to higher rightward asymmetry in inferior parietal cortex (#2R; *r =* .48 [LCBC]), leftward anterior cingulate asymmetry (ACC; #3L) was related to higher rightward asymmetry in mPFC (#6R, *r* = .47), and leftward asymmetry in a superior frontal cluster (#7L) was related to rightward asymmetry in the cingulate (#3R, *r* = .68). None of the relationships could be explained by brain size, as additionally removing the effect of intracranial volume (ICV) from cluster AI’s had a negligible effect on their interrelations (max correlation change = 0.008). Post-hoc tests confirmed that opposite-direction asymmetries were more correlated if closer in cortex (Methods); geodesic distance was lower between cluster-pairs that were more correlated (*rho* = -.35, *p* = .01 [LCBC]; -.38, *p* = .007 [UKB; Figure 5C]; -.34, *p* = .02 [HCP]), though this was driven by the aforementioned “hotspots”. By contrast, same-direction areal asymmetries were not more correlated if closer in cortex (leftward [all *p* > .5]; rightward [all *p* > .5]). This suggests specific areal asymmetries that are closer in cortex and opposite in direction may show coordinated development.

**Figure 5:**
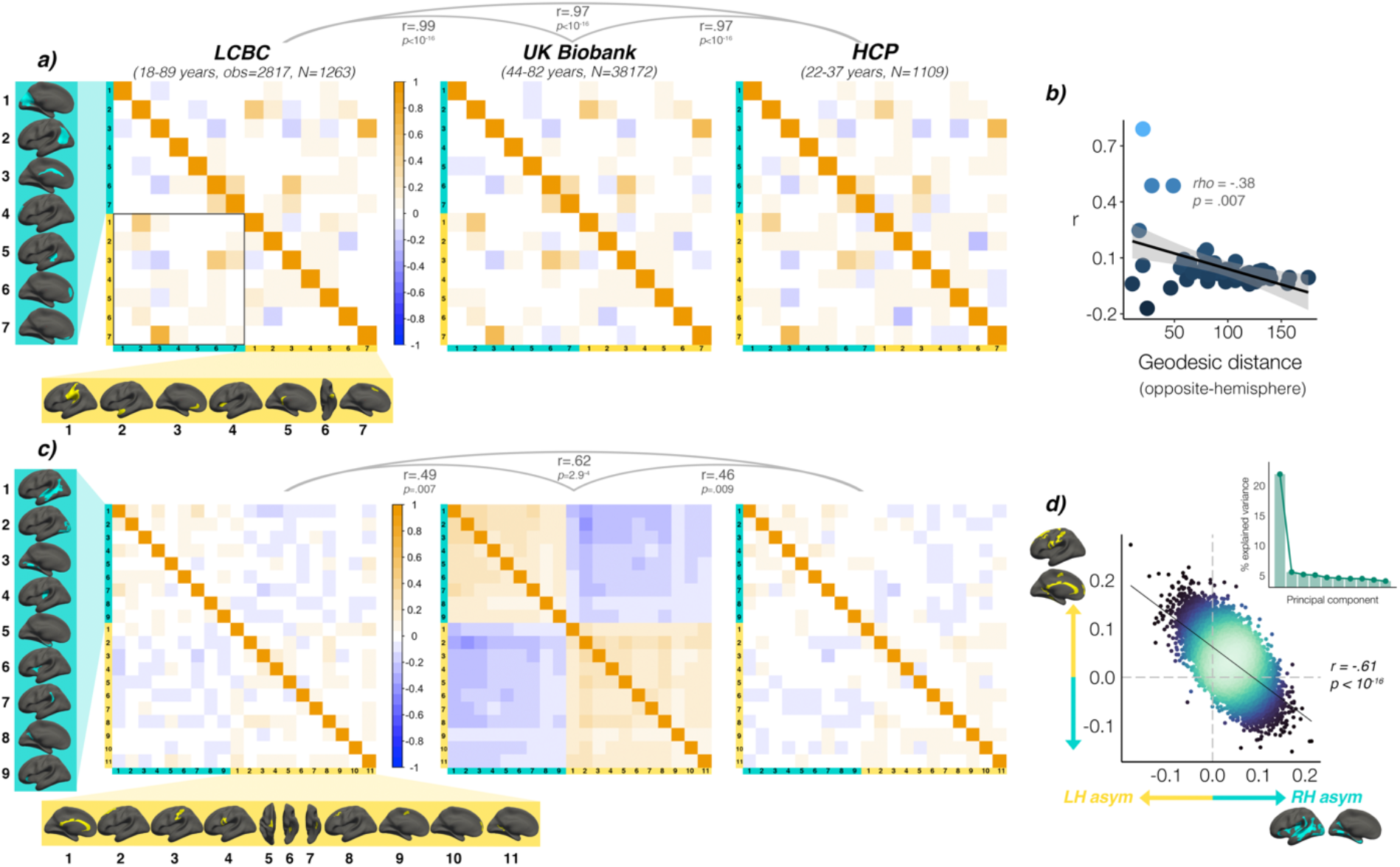
Interregional correlations between **A)** areal asymmetries and **B)** thickness asymmetries for each replication dataset (AI’s residualized for age, sex, scanner). AI’s in rightward clusters are inversed, such that positive correlations denote positive asymmetry-asymmetry relationships, regardless of direction of mean asymmetry in the cluster (i.e. higher asymmetry in the population-direction). Yellow and blue brain clusters/colours denote leftward and rightward asymmetries, respectively (clusters numbered for reference). A consistent covariance structure was evident both for areal (r >= .97) and thickness asymmetry (r >=.49; results above matrices). Black box in A highlights relationships between opposite-direction asymmetries (i.e. leftward vs rightward regions). **C)** For areal asymmetry, asymmetry in opposite-direction cluster-pairs that were closer in cortex were more positively correlated (datapoints show cluster-pairs). **D)** A single component explained 21.9% variance in thickness asymmetry in UKB (inset plot). Accordingly, we found a correlation of r = -.61 (p < 10^−16^) in UKB between mean asymmetry across leftward clusters (Y-axis) vs. mean asymmetry across rightward clusters (X-axis; AI’s in rightward clusters inversed). Lines of symmetry (0) are in dotted grey (see also Figure 4–figure supplements 1-3).

For thickness asymmetry, the correlation matrix exhibited a clear pattern in UKB that was less visible but still apparent in LCBC and HCP (Figure 5B; Figure 5–figure supplement 1B). Mantel tests confirmed that the covariance structure replicated between all dataset-pairs (LCBC-UKB *r* = .49, *p* = .007; LCBC-HCP *r* = .62, *p* < 2.9^−4^; UKB-HCP *r* = .46, *p* = .009). The observed pattern suggested higher leftward asymmetry in regions that are on average left-asymmetric was associated with less rightward asymmetry in regions that are on average right-asymmetric. However, given that the AI measure is bidirectional, closer inspection of the correlations revealed that higher leftward asymmetry in regions that are left-asymmetric actually corresponded to more *leftward* asymmetry in right-asymmetric regions, and vice versa (and on average; see Figure 5–figure supplement 2). In other words, individuals may tend towards either leftward lateralization or rightward lateralization (or symmetry) on average, irrespective of the region-specific direction of mean thickness asymmetry in the cluster. Similarly, asymmetry in left-asymmetric regions was mostly positively correlated, and asymmetry in right-asymmetric regions was mostly positively correlated. Again, additionally removing ICV-associated variance had negligible effect (max correlation change = 0.004). Post-hoc principal components analysis (PCA) in UKB revealed PC1 explained 21.9% of the variance in thickness asymmetry and suggested a single component may be evident for thickness asymmetry (Figure 5D). Accordingly, we found a correlation of r = -.61 between mean asymmetry across all leftward vs. mean asymmetry across all rightward clusters in UKB (*p* < 2.2^−16 [means weighted by cluster size]^; see Figure 4D; r = -.56, *p* < 2.2^−16 [unweighted means]^; r = .66, *p* < 2.2^−16 [PC1 across all leftward vs. PC1 across all rightward]^. Though less strong, all relationships were significant in LCBC (r = -.12; *p* = 7.4^−11 [weighted];^ r = -.05; *p* = .005 ^[unweighted]^; r = .19; *p* < 2.2^−16 [PC1 vs. PC1]^ and significant or trend-level in HCP (r = -.11; *p* = 1.6^−4 [weighted];^ r = -.04; *p* = .15 ^[unweighted]^; r = .12, *p* = 3.33^−5 [PC1 vs. PC1]^; see Figure 5–figure supplement 3). These results suggest thickness asymmetry may be globally interrelated across the cortex and show high directional variability in the adult population.

### 2.4 Heritability

Although heritability of the global measures for each hemisphere was high (area h^2^_twin_ = 92%; h^2^_SNP_ = 67%; thickness h^2^_twin_ = 81%; h^2^_SNP_ ∼ 36%), heritability of global asymmetry measures was substantially lower (area h^2^_twin_ = 16% [95% CI = 5 - 27%]; h^2^_SNP_ = 7% [3 - 10%]; thickness h^2^_twin_ = 16% [5 - 27%]; h^2^_SNP_ ∼ 1% [0 – 5%]. All estimates for global measures were post-corrected significant (*p* < .008; see Supplementary file 1G) except the SNP-based estimate for global thickness asymmetry in UKB (*p* = .22). Of our robust asymmetry clusters, four areal asymmetries showed significant twin-based heritability in HCP (post-correction; leftward SMG/perisylvian and anterior insula, and rightward cingulate and STS), all of which were also significant in the SNP-based analyses (h^2^_twin_ range = 19 – 27%; h^2^_SNP_ range = 8 – 19%). Additionally, SNP-based analyses revealed 9/14 (64%) areal asymmetry clusters exhibited significant heritability (post-correction; Supplementary file 1H). Importantly, highest SNP-based heritability was observed for leftward areal asymmetry in the anterior insula cluster (h^2^_SNP_ = 18.6%, *p* < 10^−10^; see Supplementary file 1H), which was substantially higher than the next highest estimates in SMG/perisylvian (h^2^_SNP_ = 10.7%, *p* = 3.01^−9^), retrosplenial cortex, gyrus rectus and the cingulate (all h^2^_SNP_ = 8-10%). For thickness, two robust asymmetries survived correction in the twin-based analyses (rightward STS and lingual gyrus). Of these, only STS was also significant in the SNP-based analysis but did not survive correction. Moreover, only 3/20 (15%) thickness asymmetries exhibited significant SNP-based heritability (h^2^_SNP_ = 3-7%). Cluster-wise heritability estimates were significantly lower for thickness asymmetry than for area using a SNP-based approach (*β* = -1.1, *p* = .0008; Supplementary file 1I) but not a twin-based approach (*β* = -.03, *p* = .33).

We then estimated asymmetry heritability cortex-wide ^50^ (Figure 6). For areal asymmetry, 69 parcels survived FDR-correction in HCP, 84% (58) of which were also FDR-corrected significant in the SNP-based analysis. Moreover, a total of 267 (53%) parcels exhibited significant FDR-corrected SNP-based heritability for areal asymmetry in UKB data (significant *p[FDR]*<.05 parcels in each sample are depicted with black outlines in Figure 6A). Beyond significance, a consistent heritability pattern for areal asymmetry was clearly evident using twin- and SNP-based data from independent samples, with higher heritability notably in anterior insula, SMG, Sylvian fissure, STS, calcarine sulcus, cingulate, medial and orbitofrontal cortex, and fusiform. This overlap was substantiated by a spatial correlation of r = .46 between maps; *p* < 10^−16^. Moreover, maximum SNP-heritability (cyan parcel in Figure 6) was observed in anterior insula (parcel h^2^_SNP_ = 16.4%; *p* < 10^−10^), confirming this region constitutes the most heritable cortical asymmetry in humans (and not improving on the cluster-wise estimate).

**Figure 6.**
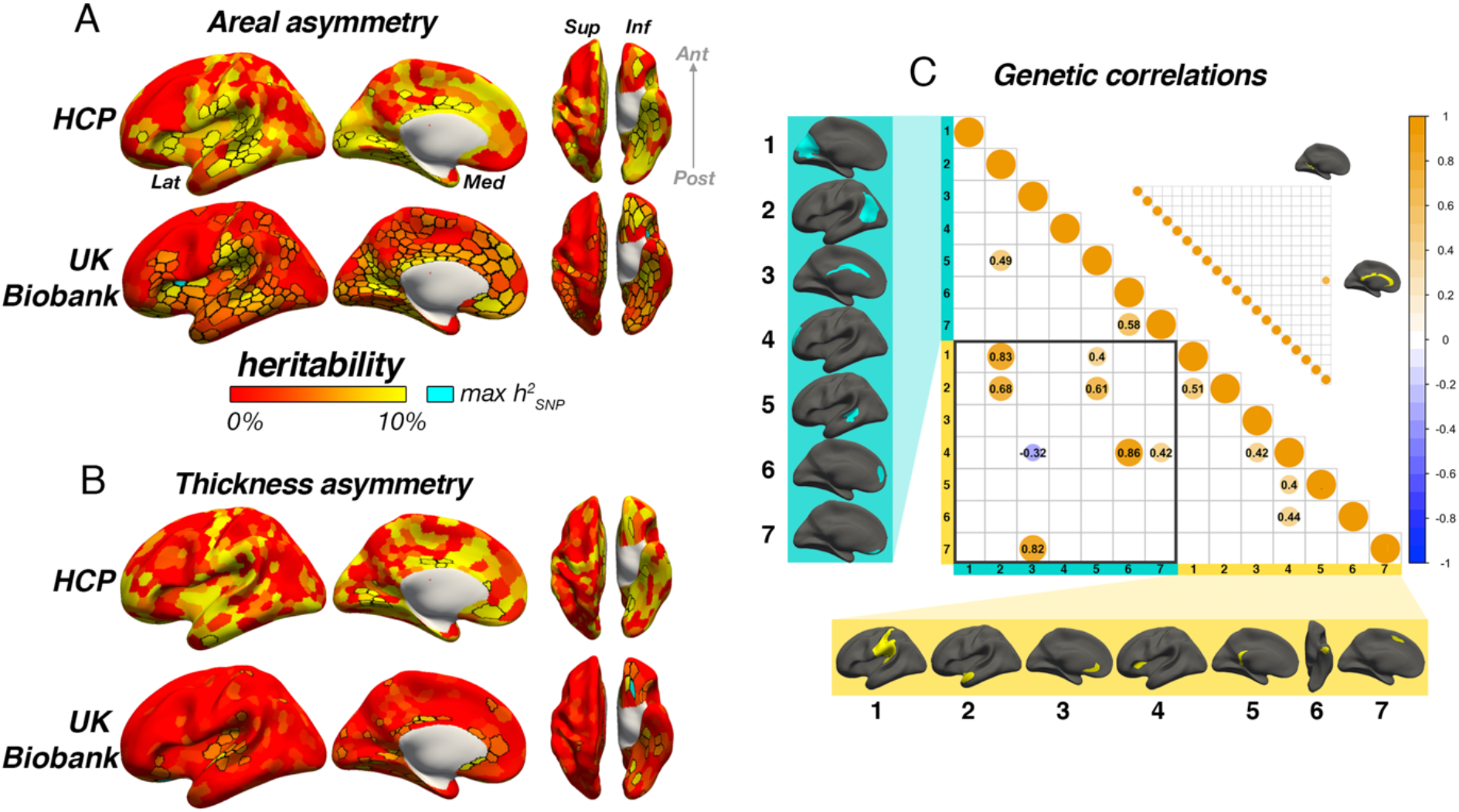
Heritability of areal **(A)** and thickness asymmetry **(B)** estimated cortex-wide using twin-based (HCP data; top row) and SNP-based methods (UKB data; bottom row). Unthresholded effect maps are shown. Parcels in black outline show significance at p[FDR]<.05. Cyan parcels depict the point of maximum observed SNP-heritability (area h^2^ = 16.4%; thickness h^2^ = 16.6%). **C)** SNP-based genetic correlations between areal (lower matrix) and thickness asymmetries (upper matrix). For area, genetic correlations explained several phenotypic correlations (Figure 5A). For thickness, one pair survived FDR-correction (shown). AI’s in rightward clusters are inversed such that positive genetic correlations denote asymmetry-asymmetry genetic relationships, regardless of direction of mean asymmetry in the cluster (i.e. higher asymmetry in the population-direction). Yellow and blue brain clusters/colours denote leftward and rightward asymmetries, respectively (clusters numbered for reference).

For thickness asymmetry, 15 parcels survived FDR-correction in the twin-based analysis, 7 of which were also post-corrected significant in the SNP-based analysis. However, significant FDR-corrected SNP-heritability was observed in only 11% (57) of parcels, including around superior temporal gyrus, planum temporale, the posterior insula/Sylvian fissure, anterior insula, and in orbitofrontal cortex (max h^2^_SNP_ = 16.6%), along the cingulate and in medial visual cortex. Beyond significance, we observed little obvious visual overlap in twin- and SNP-based heritability patterns (spatial correlation was significant but low; r = .24; *p* = .01), and higher estimates pertained to regions that were limited in extent but generally showed no clear global pattern common to both datasets, with the possible exception of calcarine and cingulate cortex. Furthermore, cortex-wide heritability estimates were significantly lower for thickness asymmetry than for area using both a SNP-based (*β* = -0.71, *p* < 2^−16^) and twin-based approach (*β* = -.33, *p* = 1.08^−7^).

For areal asymmetry, large genetic correlations explained several phenotypic correlations evident in Figure 5A (Figure 6C): high genetic correlations were found between leftward asymmetry in SMG/perisylvian and higher rightward asymmetry in lateral parietal cortex (LPC; *r*_*G*_ = .83; *p(FDR)* = <u>6.58</u>^−5^), between leftward superior frontal cortex asymmetry and rightward asymmetry along the cingulate (*r*_*G*_ = .82; *p[FDR]* = 1.15^−2^), and between leftward anterior temporal/parahippocampal asymmetry and rightward asymmetry in LPC (*r*_*G*_ = .68; *p[FDR]* = 1.16e^−2^). Genetic correlations between anterior insula and two rightward superior frontal clusters were also observed (*rG* = .86; *p[FDR]* = 1.21^−6^; rG = 0.42; *p[FDR]* = 6.64^−4^) in the absence of phenotypic correlations (Figure 5), and several same-direction asymmetries showed moderate genetic correlation. In contrast, only one post-corrected significant genetic correlation was observed for thickness asymmetry (see Figure 6C; *rG* = 0.68; *p[FDR]* = 2.97^−2^).

### 2.5 Associations with Cognition, Handedness, Sex, and ICV

Finally, several significant associations were observed between factors-of-interest and asymmetry in our clusters (Figure 7; Supplementary file 1J-K). Notably, all effect sizes were small. For general cognitive ability, we found one association, wherein higher areal asymmetry in the largest leftward cluster (SMG/perisylvian) was significantly associated with better cognition (*β* =.03 [CI = 0.02 – 0.04], *p* = 7.4^−7^, Figure 7C). Although small, we note this association was far from only just surviving correction at our predefined alpha level (*α* = .01; Methods). This was checked in the substantially reduced non-imputed subset of data with no missing cognitive variables and retained the lowest *p*-value (N = 4696; *β* =0.04 [CI = 0.01 - 0.07]; *p* = 6.9^−3^). We also post-hoc tested areal asymmetry associations with the 11 separate cognitive tests (Figure 7– figure supplement 1). For handedness, reduced leftward areal asymmetry in anterior insula and thickness asymmetry along postcentral gyrus was evident in left-handers, in line with our recent vertex-wise mapping in UKB ^22^. For sex effects, which were also small, males typically exhibited slightly stronger areal asymmetry in large clusters (e.g. leftward SMG/perisylvian and temporal pole; rightward inferior parietal and superior frontal) but reduced leftward and rightward asymmetry in mPFC. For thickness, males exhibited more rightward asymmetry in STS and posterior insula, more leftward thickness asymmetry in superior frontal cortex, but reduced rightward thickness asymmetry in entorhinal cortex and anterior insula, and reduced leftward asymmetry in caudal superior frontal cortex. As ICV effects were typically most nominal, these are shown in Figure 7–figure supplement 2. Of note, none of the reported associations changed appreciably when controlling for additional brain-size related covariates ^28^ (Supplementary file 1J-K).

**Figure 7.**
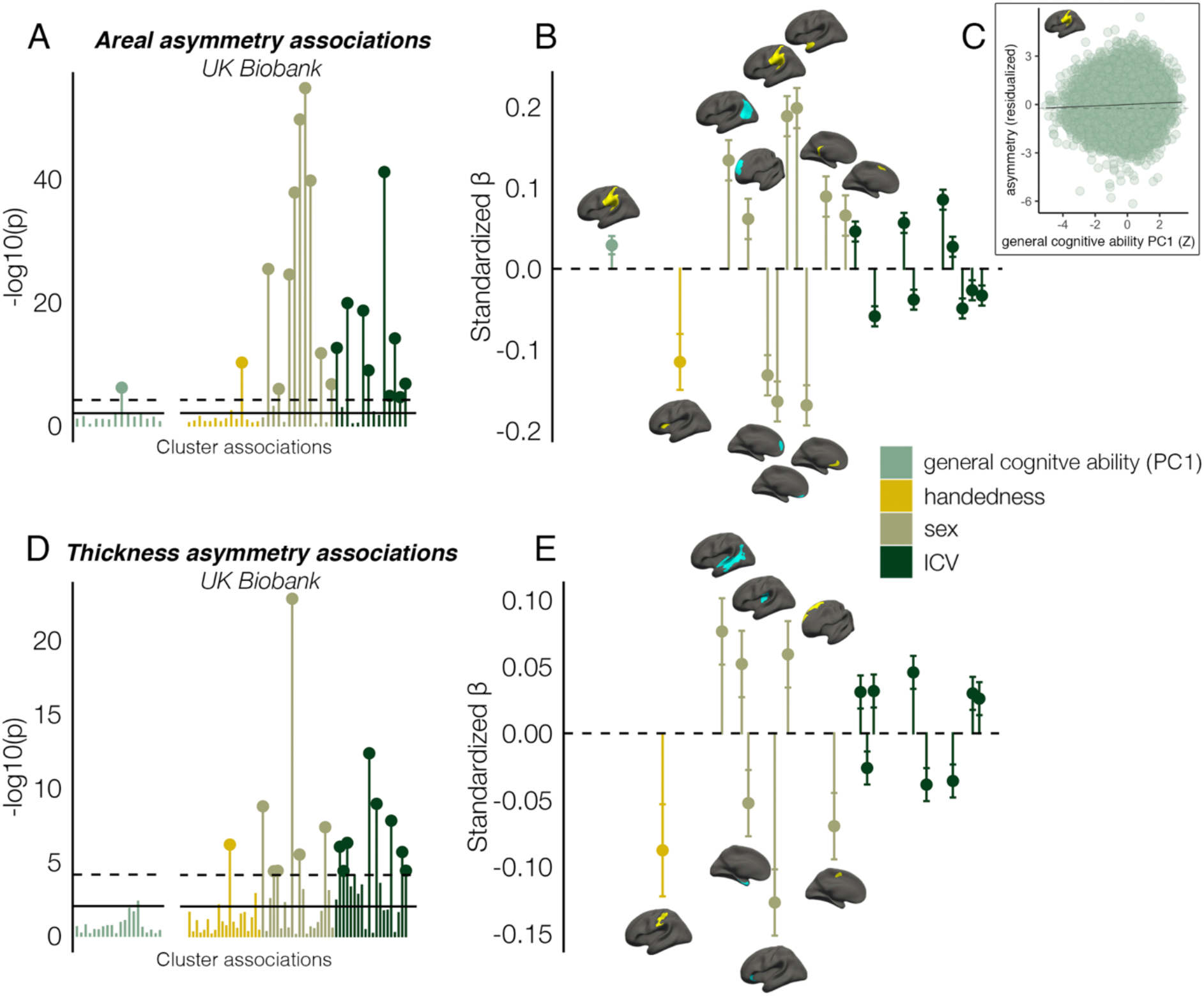
Associations with General cognitive ability (first principal component), Handedness, Sex, and Intracranial volume (ICV) in UKB, in clusters exhibiting population-level areal (upper) and thickness asymmetry (lower). **A, D)** Significance of associations (negative logarithm; corrected [p < 7.3e^−5^] and uncorrected threshold [p = .01] shown by dotted and non-dotted line, respectively). X-axis displays the test for each cluster-association. As maximum sample size was used to test each association, effects on general cognitive ability were tested in separate models with fewer observations (N = 35,199; separated association plots) than Handedness, Sex and ICV (N=37,570). C) Visualization of the found association between leftward areal asymmetry in the large supramarginal cluster upon general cognitive ability (PC1 across 11 cognitive tests). The line of null association is shown for comparison (dotted) **B, E)** Right plots denote effect sizes, 95% confidence intervals (error bars) and cortical location of associations surpassing Bonferroni-corrected significance. AIs in rightward clusters were inversed. Right handers and females are coded 0, such that a negative effect for general cognitive ability / handedness / sex / ICV / denotes less asymmetry in higher cognition / left handers / males / larger brains. Associations with ICV are shown in Figure 7–figure supplement 2. Yellow and blue clusters denote leftward and rightward asymmetries, respectively.

## 3. Discussion

Combining the strengths of a vertex-wise delineation of population-level cortical asymmetry in 7 international datasets and dense longitudinal data, we offer the first description of the longitudinal developmental and lifespan trajectories of cortical asymmetry, advancing knowledge on normal human brain development. We show areal asymmetry is predominantly stable across life, whereas we trace developmental growth in many thickness asymmetries signifying differentiation of thickness asymmetry from early childhood to the mid-20s. We further demonstrate the replicable interregional relationships between asymmetries within individuals, provide the most detailed heritability maps for cortical asymmetry to date, and uncover novel and confirm previously-reported associations with factors purportedly important in the context of asymmetry – all with small effects. All maps are available at neurovault.org/XXXX.

### 3.1. Population-level asymmetry

Our vertex-wise analysis of cortical asymmetries that reproduce across adult cohorts replicates and completes a recent low-resolution meta-analysis ^7^, and can serve as a high-fidelity phenotype for future brain asymmetry studies. The marked consistency across samples here suggests consensus may now be reached regarding cortical asymmetry phenotypes in humans, as our results agree with most reported results for areal asymmetry ^7,8,11,21,31^, as well as several, typically more recent reports for thickness asymmetry ^7,17–19^. Indeed, for thickness asymmetry – for which findings have been particularly mixed ^7,8,18–20,10–17^ – the left-right patterning here is compatible with low-resolution meta-analyses ^7^, asymmetries evident in the first months of life ^19^, a large-scale mapping in mid-old age ^22^, reports using alternative analysis streams ^17,19^, and possibly the overall pattern of brain torque ^7,51^ - a gross hemispheric twist leading to frontal and occipital bending at the poles ^52^. The high overlap in effects between 7 datasets from 4 countries here suggests these results likely apply universally. This evident consensus suggests genetic-developmental programs regulate mean brain lateralization with respect to both area and thickness in humans. However, the genetic findings presented herein suggest these may have reached population fixation, as heritability of even our optimally delineated asymmetry measures was generally low. This indicates either subject-specific stochastic mechanisms in early neurodevelopment or later developmental influences primarily determine cortical asymmetry. Tracing their lifespan development, we show the lifespan trajectories of areal asymmetry primarily suggest this form of cerebral asymmetry is developmentally stable at least from age ∼4, maintained throughout life, and formed early on – possibly *in utero* ^27,31^. One interpretation of lifespan stability combined with low heritability may be stochastic early life developmental influences determine interindividual differences in areal asymmetry more than later developmental change. However, future work linking prenatal and childhood trajectories is needed to affirm this. Still, we also found relatively stronger heritability and reproducible heritability maps for areal asymmetry (notably, anterior insula exhibited ∼19% SNP-based heritability). This also illustrates region-dependent interindividual genetic effects upon areal asymmetry, and the fact that phenotypic correlations were underpinned by high genetic correlations suggests specific areal asymmetries are formed under common genetic influence. In contrast, our findings of childhood development of thickness asymmetry until a peak around age ∼24 ^18^, higher directional variability in adult samples, and lower heritability, converge to suggest thickness asymmetry may be more shaped through subject-specific effects in later childhood development, possibly as the brain grows in interaction with the environment. This interpretation applied to asymmetry agrees with work suggesting cortical area in general may trace more to early life factors ^53–55^ whereas thickness may be more related to and impacted by lifespan influences ^54,56^.

### 3.2. Lifespan trajectories

Our longitudinal description of cortical asymmetry lifespan trajectories gleaned novel insight into normal brain maturation and development. For areal asymmetry, adult-like patterns of lateralization were strongly established already before ∼4 years, indicating areal asymmetry traces back further and does not primarily emerge through later cortical expansion ^57^. Rather, the lifespan trajectories predominantly show stability from childhood to old age, as areal asymmetry was generally maintained through periods of developmental expansion and aging-associated change that were region-specific and bilateral. This agrees with evidence indicating areal asymmetry may be primarily determined *in utero* ^31^, and indirect evidence suggesting little change in areal asymmetry from birth to 2 years despite rapid and concurrent cortical expansion ^31,32,57^. It may also fit with the principle that the primary microstructural basis of cortical area ^55^ – the number of and spacing between cortical minicolumns – is determined in prenatal life ^55,56^, and agree with evidence suggesting asymmetry at this microstructural level may underly hemispheric differences in surface area ^58^. The developmental trajectories agree with studies indicating areal asymmetry is established and strongly directional early in life ^30,31^. That anatomical change in later development specifically in surface area follows embryonic gene expression gradients may also agree with a prenatal account for areal asymmetry ^56^. Our results may therefore constrain the extent to which areal asymmetry can be viewed as a plastic feature of brain organization, and may even suggest areal asymmetry may sometimes be a marker for innate hemispheric specializations shared by most humans. Although future research is needed to characterize asymmetry structure-function relationships, the high degree of precision with which a large leftward areal asymmetry follows the contours of auditory-related regions in the Sylvian fissure (Figure 1–figure supplement 1) which show left functional lateralization in humans may be one example ^58–60^; we found ∼94% of individuals exhibited leftward areal asymmetry here – the most consistently lateralized cortical region in humans (Figure 1F).

In stark contrast, although weak thickness asymmetry was evident by age 4, we observed considerable childhood developmental growth in many regional thickness asymmetries. Developmental trajectories showed non-linear asymmetry growth by virtue of accelerated thinning of the non-dominant hemisphere (10/20 clusters showed a relative hemispheric difference in the rate of developmental thinning; Figure 4). This led to maximally established asymmetry around ∼24 years of age. These trajectories clearly suggest differentiation of the cortex is occurring with respect to thickness asymmetry in development, possibly (though not necessarily) suggesting thickness asymmetry may be more amenable to experience-dependent change. Indeed, as cortical thinning in childhood is thought to partly reflect likely learning-dependent processes such as intracortical myelination ^61^ and possibly pruning of initially overproduced synapses ^62,63^ and neuropil reduction, thickness asymmetry growth may suggest hemispheric differences in the developmental optimization of cortical networks at least partly shaped by childhood experience. This raises the possibility thickness asymmetry may be a marker of ontogenetic hemispheric specialization within neurocognitive networks, and may also fit with animal models suggesting lateralized training can alter hemispheric imbalance of cortical thickness ^64^. However, this interpretation is speculative, and our results could be used to guide intervention-based approaches to experimentally test plasticity of thickness asymmetry in humans. Our lifespan results are often difficult to reconcile with earlier lifespan reports finding different mean thickness asymmetry patterns ^10,12,15^ and modelling asymmetry linearly across cross-sectional developmental and adult age-ranges ^10,15,17^. However, our findings in development agree with work finding a similar left-right thickness asymmetry pattern shows rapid longitudinal asymmetry increase in the first years of life ^19^, with especially rapid increase in mPFC ^19^. As we observed rapid asymmetry differentiation that spanned across childhood and adolescence in most prefrontal regions and several others (Figure 3; Figure 4; Figure 3–figure supplements 3 & 4), we extend these earlier findings in neonates ^19^. Likely, the combination of longitudinal data and nonlinear modelling was critical to capture early developmental growth in thickness asymmetry within a lifespan perspective, because large interindividual variation in hemispheric thickness estimates at any age may hinder detection of subtle change effects even in large cross-sectional samples ^65^, possibly rendering follow-up data all but essential in the context of cortical asymmetry change. As prefrontal thickness asymmetry seems particularly vulnerable in some neurodevelopmental disorders ^24^, aging, and Alzheimer’s disease ^18^, these trajectories may provide a useful normative reference, for example regarding the timing of expected brain change in development. With regards to aging, most clusters delineated here did not exhibit the relative hemispheric difference in rates of cortical thinning we have previously shown is a feature of aging in heteromodal cortex ^18^, except in clusters overlapping with our previous analysis (Figure 3– figure supplement 5). This agrees with our previous work showing aging-related loss in thickness asymmetry is specific to heteromodal cortical regions vulnerable in aging. In these regions, we also found strong evidence of early developmental growth in thickness asymmetry (Figure 3–figure supplement 4). Developmental differentiation and aging-related dedifferentiation of thickness asymmetry underscores its proposed role in supporting optimal brain organization and function, though it seems not all thickness asymmetries that grow in childhood development decline in aging.

### 3.3. Interregional correlations

For areal asymmetry, we uncovered a covariance structure that almost perfectly replicated across datasets. In general, this fit with a multifaceted view ^29,34,35^, in which most asymmetries were either not or only weakly correlated – but reliably so – contrasting views emphasizing a single biological ^37,38^ or overall anatomical factor ^66^ controlling cerebral lateralization. However, we also identified several regions wherein areal asymmetry reliably correlated within individuals, showing the variance in cortical asymmetries is not always dissociable, as often thought ^29,34,35^. The strongest relationships all pertained to asymmetries that were proximal in cortex but opposite in direction. Several of these were underpinned by high asymmetry-asymmetry genetic correlations, illustrating cerebral lateralizations in surface area that are formed under common genetic influence, possibly in agreement with likely prenatal origins for areal asymmetry ^31,55^.

For thickness asymmetry, we also uncovered a common covariance structure – particularly clear in UKB – that nevertheless replicated with moderate precision across datasets (Figure 5C). Furthermore, a single component explained a relatively high proportion of variance in thickness asymmetry in UKB, and a high correlation across 38,172 individuals further suggested thickness asymmetry may be globally interrelated across the cortex (Figure 5D). These data for thickness indicate individuals may tend towards either leftward asymmetry, rightward asymmetry, or symmetry, both globally across the cortex and irrespective of the region-specific average direction of asymmetry (Figure 5–figure supplements 2-3). Though it is unclear why the relationships were weaker in the other datasets tested, we nevertheless found similarly significant relationships in each (Figure 5–figure supplement 3). This result seems in broad agreement with the notion that some lateralized genetic-developmental programs may trigger lateralization in either direction ^35^ or lose their directional bias through environmental interaction ^35^. As thickness asymmetry seems established at but minimal from birth ^19^, genetic effects may determine the average region-specific hemispheric bias in the population, but later developmental change may subsequently confer major increases upon its directional variance ^35^. Overall, the evidence converges to suggest a high degree of developmental change may shape thickness asymmetry and lead to higher directional variability in the population. Thus, far from being independent phenotypes ^29,34^, thickness asymmetries may be globally interrelated across the cortex and their direction coordinated through development.

### 3.4. Genetic influences

For areal asymmetry, we found replicable patterns of low-moderate heritability across datasets and across twin and genomic methods. We also found areal asymmetry in the anterior insula is, to our knowledge, the most heritable brain or behavioural asymmetry yet reported with genomic methods ^22,27,67–69^, with common SNPs explaining ∼19% variance. This is a substantial improvement on our recent report of < 5% ^22^, and illustrates a benefit of our data-driven population-mapping approach. As we reported recently ^22^, we confirm asymmetry in this region associates with handedness. Furthermore, highest SNP and twin-based heritability for areal asymmetry was found in all regions that constitute the earliest emerging cortical asymmetries *in utero* ^70–73^: anterior insula, STS, PT, medial occipital cortex, and parahippocampal gyrus (Figure 6A). However, observations of significant heritability were not restricted to these regions, as most areal asymmetries exhibited significant – albeit often lower – SNP-based heritability, as did most parcels when estimated cortex-wide, and significant SNP-based heritability was also evident in regions not found in the present analyses to show strong areal asymmetry, such as Broca’s area (but see Figure 1–figure supplement 4). These effects agree with and elaborate on two genetic explorations using atlas-based methods ^7,27^ and reports of heritable areal asymmetry in handedness-associated clusters ^22^. However, as noted above, although heritability of either hemisphere was high, areal asymmetry heritability was still only moderate at best, suggesting both genetics but primarily subject-specific stochastic effects likely underly its formation. By contrast, thickness asymmetry was generally not heritable, or showed low and localized heritability effects with no clear global pattern. We also observed more divergent results using twin and genomic methods for thickness, possibly due to low-power for twin-models, though we note the SNP-based effects we observed were somewhat in agreement with a previous twin study ^7^.

Considered together, lifespan stability possibly from birth ^31^, less interindividual directional variability, higher heritability, and phenotypic and genetic correlations all converge to suggest comparatively higher genetic influence upon interindividual differences in areal asymmetry. This also agrees with work showing genetic variants associated with (mostly areal) asymmetry are primarily expressed in prenatal life ^27^. By contrast, childhood developmental change, high interindividual directional variability and low heritability for thickness asymmetry converge to fit a scenario whereby thickness asymmetry may be more shaped through individual lifespan exposures ^56^, or where its developmental direction may be triggered by random influences ^74^. Whether region-specific thickness asymmetry change relates to the maturation of lateralized brain functions ^74,75^ will be an important question for future research. Regardless, our results support a differentiation between early-life and later developmental factors in shaping areal and thickness asymmetry, respectively.

### 3.5. Individual differences

Screening population-level asymmetries for association with general cognitive ability revealed one region – SMG/perisylvian – wherein higher leftward areal asymmetry related to higher cognition. Interestingly, this cluster was consistently the most lateralized across individuals, with ∼94% directional concordance (Figure 1G), suggesting highly regulated genetic-developmental programs shape its laterality in humans. Asymmetry here is likely related to brain torque, which leads to interhemispheric anatomical differences especially around the Sylvian fissure ^52,76^. This may therefore agree with recent work suggesting brain torque relates to cognitive outcomes ^77,78^. However, as seems typical of brain associations in big data ^79^ and may be expected for any single structural measure explaining a complex phenomenon, the association was notably small. Also of note, no other asymmetry showed evidence of cognitive association (Figure 7–figure supplement 1), suggesting previously reported associations in small samples will likely not replicate ^17^. That the association we find was specific to the most lateralized areal asymmetry in humans may suggest disruptions in early life cerebral lateralization lead to cognitive deficits – detectable in later life as small effects in big data. This may also agree with work suggesting differences in general cognitive ability relate primarily to areal phenotypes ^53,56^ formed early in life ^55^.

Consistent with our recent vertex-wise analysis in UKB ^22^, we confirmed leftward areal asymmetry of anterior insula, and leftward somatosensory thickness asymmetry is subtly reduced in left handers. Sha et al. ^22^ reported shared genetic influences upon handedness and cortical asymmetry in anterior insula and other more focal regions not identified here. Anterior insula lies within a left-lateralized functional language network ^80^, and its structural asymmetry may relate to language lateralization ^42,81,82^ in which left-handers show increased incidence of atypicality ^48,83,84^. Since asymmetry here emerges early *in utero* ^70^ and is by far the most heritable (see above), we agree with others ^42^ that future research will find this ontogenetically foundational region of cortex ^85,86^ fruitful in understanding genetic-developmental mechanisms influencing laterality. Leftward thickness asymmetry reduction in somatosensory cortex in left handers also echoes our recent report ^22^, and may also fit a scenario whereby thickness asymmetries are partly shaped through use-dependent plasticity, possibly carrying information regarding group-level hemispheric specializations of function. However, the small effects highlight that cortical asymmetry cannot predict individual hand preference. Similarly, asymmetry-relationships with other factors typically assumed important were all small – despite our optimization of asymmetry phenotypes – and mostly compatible with those reported in the ENIGMA meta-analysis ^7^ and elsewhere ^28,41^. Concerning sex effects – which were small, differing in direction, and more predictive than ICV ^41^ – inconsistencies evident between ours and ENIGMA include findings of increased (here) and decreased ^7^ lateral parietal areal asymmetry in males, and increased ^7^ and decreased (here) entorhinal thickness asymmetry in males, and our approach detected other regions slightly more asymmetric in males (e.g. STS). Possibly, differences in sample median age (here UKB = ∼64; Kong et al. = 26 ^7^) and potential sex-differences in age decline trajectories ^87^ may underlie some inconsistencies, maybe more so for thickness measures in structures vulnerable to age-related degeneration ^22^.

### 3.6. Limitations

Several limitations should be mentioned. First, our delineation of population-level asymmetry used a single analysis software. As with most current papers we used FreeSurfer’s default ‘recon-all’ function to delineate the cortex, which has been extensively validated against postmortem measurements ^88^ and is the software underlying most large-scale studies involving brain measures. It is currently unclear to what extent differences in pipelines account for previous mixed results ^7,8,18–20,10–17^. Although we highlight there are clear commonalities between our results and studies using alternative pipelines ^17,19,20^, which may suggest our results generalize across analysis systems ^17,19–21^, we found one instance where the MRI pipeline leads to different results for thickness asymmetry (Figure 1–figure supplement 4). It is not known what underlies this difference, though we find it is unrelated to the cross-hemispheric registration methods employed here, as our results reproduce using standard parcellation methods and thus are likely evident in most FreeSurfer-derived datasets (Figure 1–figure supplement 5). One possibility could be that thickness asymmetry may not reflect cortical thickness differences per se, but rather reflect biologically meaningful hemispheric differences in intracortical myelination levels that are consistently picked up on via FreeSurfer’s standard delineation procedure. That the thickness asymmetry pattern we observe shows a clear developmental trajectory also suggests it is a true biological effect (Figure 3; Figure 4; Figure 3–figure supplements 1-4). Conceivable sources of inconsistently reported results for cortical asymmetries likely include varying age-distributions (particularly for thickness) ^18^, and multiple asymmetries within atlas-based parcels (e.g. we observed notably discrepant results to ENIGMA for insula thickness asymmetry ^7^). Still, this does not explain other discrepant reports, such as areal asymmetry of STS, which was rightward in all samples here (Figure 1–figure supplement 2; cf. ^7,29,30^). Thickness asymmetries seem also more variable compared with area ^7^, though it should be noted thickness in general may be slightly less reliable ^89^, the asymmetry effect is smaller, and thus possibly contains more error. Relatedly, while the reported areal and thickness asymmetry patterns and strengths using cross-hemispheric methods agree with standard analysis (Figure 1–figure supplement 5), the magnitude of some asymmetries near the boundary of the subcortex may be exaggerated via this approach ^22^. And although we did not find (strong) areal asymmetry in inferior frontal regions as reported by Kong et al ^7^, both the unthresholded significance maps and standard parcellation analyses were compatible with this (Figure 1–figure supplement 2 & 5), highlighting a limitation of our chosen method. Second, while GAMMs are considered an optimal modelling technique for longitudinal lifespan data and are robust to nonuniform age distributions ^90^, relative underrepresentation of the mid-adulthood age-range may drive trajectory inflection points around this age ^18^, suggesting caution is warranted regarding interpreting mid-life inflection points as reflecting real change. Relatedly, as individual-level mean estimates likely contain more measurement error when extracted from smaller clusters, the lifespan trajectories from smaller clusters may be more variable, complicated, and prone to deflection by density differences along the age distribution. As can be seen in Figures 3-4, spatially averaging across asymmetries helps smooth out some of this noise to better reveal developmental principles underlying structural asymmetries, but also trades off with accuracy, since the trajectories from distinct regions do somewhat differ. Third, though the differing heritability methods enabled a replication test, important differences between methods should be considered when interpreting heritability estimates, and may partly explain the common observation of higher estimates from twin methods, which we also observed. For example, twin methods implicitly incorporate additive genetic effects and gene-gene interactions amongst other terms, whereas SNP-based heritability incorporates only additive genetic effects. Still, twin methods may in some cases be prone to overestimating heritability due to unmet assumptions, particularly where the shared environment is thought to play a role ^91^, whereas SNP-based methods may not capture all phenotype-relevant genetic variance and have their own assumptions ^92^. We also observed that twin-based estimates were often substantially higher also where non-significant, which may suggest low power to detect the effects in question in HCP family data, or may agree with calls that caution against overinterpreting twin-based estimates ^91^. Adding to this complexity, the samples used for twin- and SNP-based estimation consist of young adults and older adults, respectively. Hence, it is possible low SNP-based estimates for thickness asymmetry in UKB may be partly due to reduced mean thickness asymmetry in older adults ^18,28^. However, as cortex-wide twin-based estimates for thickness asymmetry were also significantly lower than for areal asymmetry in the young HCP sample, this suggests age is likely not the main driver behind this difference, but a confound that further hinders comparison of results between methods. Again, we also cannot rule out that potential reliability differences between thickness and areal asymmetry may partially explain some magnitude differences in heritability estimates. Further, genetic correlations can be high even where heritability of either trait is low, and are typically higher than phenotypic correlations – known phenomena also observed here ^93,94^. It may therefore be prudent to not overinterpret their magnitudes, though we emphasize their veracity seems well-supported, as one would expect true genetic correlations between developmentally-related and similar traits (here, sampled from nearby in the same organ), and to track the phenotypic correlations – both of which we also observed here – indicative of synchronized development of areal asymmetries through common genetic causes. Fourth, we imposed a necessary cluster size limit for overlapping asymmetry effects across samples, and thus more focal asymmetries may also be informative in relation to the factors tested here ^22^. Fifth, as only dichotomous handedness self-reports are available with UKB, future studies might benefit from incorporating more nuanced handedness assessments not currently available in data of this size. Relatedly, because UKB cognitive data is not exhaustive ^95^ (e.g. fluid IQ ranges from 1-13), we extracted the common variance across core tests to index general cognitive ability. This approach does not permit testing associations with well-operationalized or specific cognitive domains (for which UKB cognitive data may not be sufficient), and it remains to be seen whether cortical asymmetry may be informative in the context of specific forms of lateralized cognition ^96^, or using absolute non-directional measures. However, the subtlety of the only effect on cognition we find here may suggest we might expect similarly weak effects, though not necessarily if brain and phenotypic measurement accuracy is jointly optimized ^97^.

Overall, we track the development of population-level cerebral cortical asymmetries longitudinally across life and perform analyses to trace developmental principles underlying their formation. Developmental trajectories, interregional correlations and genetic analyses converge upon a differentiation between early-life and later-developmental factors underlying the formation of areal and thickness asymmetries, respectively. By revealing hitherto unknown principles of developmental stability and change underlying diverse aspects of cortical asymmetry, we here advance knowledge of normal human brain development.

## 4. Methods

### 4.1 Samples

We used anatomical T1 -weighted (T1w) scans from 7 independent international MRI datasets originating from 4 countries (see Supplementary file 1A for an overview of samples used for each analysis). Note that with the exception of vertex-wise analyses in UKB (see below), all analyses made use of all available observations from each sample meeting the stated age-range criteria for each analysis.

#### 4.1.1 Reproducibility across samples: population-level asymmetry

To delineate average adult patterns of whole-cortical areal and thickness asymmetry, we restricted the age-range of all samples used in the vertex-wise analyses to 18-55. ***Dataset 1*:** Here, the Center for Lifespan Changes in Brain and Cognition (*LCBC*) sample comprised 1572 mixed cross-sectional and longitudinal scans (N longitudinal = 812; timepoint range = 1-6) from 923 participants (mean age = 30.6 ± 9.6) collected across 2 scanners. Additionally, 125 individuals were double-scanned at the same timepoint on both scanners. ***Dataset 2:*** The Cambridge Centre for Ageing and Neuroscience (*Cam-CAN*) ^98^ sample comprised cross-sectional scans of 321 individuals (mean age = 38.7 ± 9.7) ^99^. ***Dataset 3:*** The Dallas Lifespan Brain Study (*DLBS*) ^100^ sample comprised cross-sectional scans of 160 individuals (mean age = 37.5 ± 10.7). ***Dataset 4:*** The Southwest University Adult Lifespan Dataset (*SALD*) ^101^ sample comprised crosssectional scans of 301 individuals (mean age = 33.7 ± 11.5). ***Dataset 5:*** The *IXI* sample comprised cross-sectional scans of 313 healthy individuals collected across 3 scanners (mean age = 36.8 ± 9.6; http://brain-development.org/ixi-dataset). ***Dataset 6:*** Here, the Human Connectome Project (*HCP*) 1200 ^102^ sample comprised 1111 scans (mean age = 28.8 ± 3.7). ***Dataset 7:*** Here, the UKB sample consisted of 1000 randomly sampled cross-sectional scans (mean age = 52.1 ± 1.9), restricted to be comparable in size to the other datasets in this analysis.

#### 4.1.2 Lifespan trajectories

Here, we used the full age-range of the longitudinal lifespan LCBC sample (4.1 - 89.4 years), 3937 cross-sectional and longitudinal scans (N longitudinal = 2762) from 1886 individuals (females = 1139; mean age = 36.8) collected across 4 scanners (271 double-scans) ^65,103^.

#### 4.1.3 Interregional correlations

Here, we used the three largest datasets: LCBC (N = 1263; N obs = 2817), UKB (N = 38,172), and HCP (N = 1109; two outliers removed; see 4.3.3), excluding the childhood age-range from the LCBC sample not covered in any other dataset.

#### 4.1.4 Heritability and individual differences

For twin heritability, we used HCP 1200 extended twin data (1037 scans from twins and non-twin siblings; age-range = 22-37; mean age = 28.9 ± 3.7). The various kinships are described in Supplementary file 1B. All included twin pairs were same-sex. For SNP-heritability, we used the UKB imaging sample with genome-wide data surpassing quality control (N = 31,433; see 4.3.4). For individual differences analyses, we used the UKB imaging sample with the maximum number of available observations for each variable-of-interest (see 4.3.5).

### 4.2. MRI preprocessing

T1w anatomical images (see Supplementary file 1C for MRI acquisition parameters) were processed with FreeSurfer (v6.0.0) ^104^ and vertex-wise areal and thickness morphometry estimates were obtained for each MRI observation. As the LCBC sample also contained longitudinal observations, initial cross-sectional reconstructions in LCBC were subsequently ran through FreeSurfer’s longitudinal pipeline. As HCP data was acquired at a higher voxel resolution (0.7mm isotropic), the T1w scans were processed with the --hires flag to recon-all ^105^. Areal and thickness maps of the LH and RH of each participant in each dataset were resampled from the native cortical geometry to a symmetrical surface template (“*LH_sym*”) ^11,106^ based on cross-hemispheric registration ^107^. This procedure achieves vertex-wise alignment of the data from each participant and homotopic hemisphere in a common analysis space, enabling a whole-cortical and data-driven analysis of cortical asymmetry. Areal values were resampled with an additional Jacobian correction to ensure preservation of the areal quantities ^108^. We then applied an 8mm FWHM Gaussian kernel to surface-smooth the LH and RH data.

### 4.3 Data analysis

All analyses were performed in FreeSurfer (v6.0) and R (v4.1.1).

#### 4.3.1 Population-level asymmetry

We assessed areal and thickness asymmetry vertex-wise using FreeSurfer’s Linear Mixed Effects (LME) tool ^109^. Asymmetry was delineated via the main effect of Hemisphere (controlling for Age, Age × Hemisphere, Sex, Scanner [where applicable], with a random subject term). For each sample and metric, we computed mean Asymmetry Index maps (AI; defined as (LH-RH) / ((LH+RH)/2)). Spatial overlap of AI maps across datasets was quantified by correlating the 1-dimensional surface data between every dataset pair (Pearson’s r). Next, to delineate regions exhibiting robust areal and thickness asymmetry across datasets, we thresholded and binarized the AI maps by a given absolute effect size (areal = 5%; thickness = 1%; achieving *p*[FDR] < .001 in most datasets with FreeSurfer’s 2-stage FDR-procedure ^109^), and summed the binary maps. After removing the smallest clusters (<200 mm^2^), a set of robust clusters was defined as those exhibiting overlapping effects in 6 out of 7 samples. We then extracted area and thickness data in symmetrical space for each cluster, subject, and hemisphere, spatially averaging across vertices.

#### 4.3.2 Lifespan trajectories

Factor-smooth GAMMs (“gamm4” ^110^) were used to fit a smooth Age trajectory per Hemisphere, and assess the smooth Age × Hemisphere interaction in our clusters. GAMMs incorporate both cross-sectional and longitudinal data to capture nonlinearity of the mean level trajectories across persons ^111^. The linear predictor matrix of the GAMM was used to obtain asymmetry trajectories and their confidence intervals, computed as the difference between zero-centered (i.e. demeaned) hemispheric age-trajectories. We included Hemisphere as an additional fixed effect, sex and scanner as covariates-of-no-interest, and a random subject intercept. We did not consider sex differences in lifespan asymmetry change because our elected method was not well-suited to testing three-way interactions between nonlinear smooth terms. A low number of basis dimensions for each smoothing spline was chosen to guard against overfitting (knots = 6; see Figure 3–figure supplement 1). LCBC outliers falling > 6SD from the trajectory of either hemisphere were detected and removed on a region-wise basis (Supplementary file 1E-F). To calculate relative change, we refitted lifespan GAMMs adding an ICV covariate, then scaled the LH and RH fitted lifespan trajectories by the prediction at the minimum age (i.e. ∼4 years). Age at peak thickness asymmetry was estimated where the CI’s of absolute hemispheric trajectories were maximally non-overlapping.

#### 4.3.3 Interregional correlations

We assessed covariance between asymmetries, separately for areal and thickness asymmetry. Here, we regressed out age, sex and scanner (where applicable) from each AI, using linear mixed models after collating the data from each sample (adding random intercepts for LCBC subjects), to ensure the correction was unaffected by differences in sample age-distribution. Separately for each dataset, we then obtained the cluster-cluster correlation matrix. All individual AI’s in clusters with rightward mean asymmetry were inversed, such that positive correlations denote asymmetry-asymmetry relationships regardless of the direction of mean asymmetry in the cluster (*i.e*. higher asymmetry in the population-direction). At this point, two strong outliers in HCP data were detected and discarded for this and all subsequent analyses (Figure 4–figure supplement 4). Replication was assessed using the Mantel test (“ade4” ^112^) between each dataset-pair (LCBC, UKB, HCP) across 10,000 permutations. We then post-hoc tested whether covariance between areal asymmetries was related to proximity in cortex, obtaining the average geodesic distance between all clusters along the ipsilateral surface (“SurfDist” Python package ^113^), and correlating pair-wise distance with pair-wise correlation coefficient (Fisher’s transformed coefficients; Spearman’s correlation). To post-hoc assess whether observed covariance patterns for thickness asymmetry reflected a global effect, we ran a PCA across z-transformed AI’s for all thickness clusters (pre-corrected for the same covariates). Based on the results, we computed the mean AIs across all leftward clusters, and across all rightward clusters, and tested the partial correlation between mean leftward thickness asymmetry in left-asymmetric clusters and mean rightward thickness asymmetry in right-asymmetric clusters, in each of the three cohorts.

#### 4.3.4 Heritability

Heritability of areal and thickness asymmetry was assessed using both twin- and SNP-based methods, both for our set of robust clusters and cortex-wide across 500 parcels ^50^. For cluster analyses, significance was considered at Bonferroni-corrected p<.05 applied separately across each metric. Cortex-wide significance was considered at *p(FDR)* < .05 (500 tests per map). Twin heritability was assessed using AE models in “OpenMx” ^114^, which use observed cross-twin and cross-sibling covariance to decompose the proportion of observed phenotypic variance into additive genetic effects [A], and unique environmental effects or error [E]. Data were reformatted such that rows represented family-wise observations. As is standard, we set A to be 1 for MZ twins assumed to share 100% of their segregating genes (but see ^115^), and 0.5 for DZ twins and siblings that share 50% on average. For each phenotype we first regressed out age and sex and computed z-scores. Statistical significance was assessed by comparing model fit to submodels with the A parameter set to 0.

For SNP-heritability, the final genetic sample consisted of 31,433 UKB participants (application #32048) with imaging and quality checked genetic data. We removed subjects that were outliers based on heterozygosity [field 22027] and missingness (> 0.05), mismatched genetic and reported sex [22001], sex chromosome aneuploidies [22019], and those not in the “white British ancestry” subset [22006] ^116^. At variant level, after removing SNPs with minor allele frequency < 0.01, data from 654,584 autosomal SNPs were used to compute a genetic relationship matrix using GCTA (v1.93.2) ^117^. For each phenotype, we first regressed out age and sex and computed z-scores. Genome-based restricted maximum likelihood (GREML) methods as implemented in GCTA were then used to compute SNP-heritability for each AI measure, applying a kinship coefficient cut-off of 0.025 (excluding one individual from each pair), and controlling for genetic population structure (first ten principal components). Bivariate GREML analysis was used to test genetic correlations between asymmetry clusters ^117^. These estimate the proportion of variance two asymmetries share due to genetic influences through pleiotropic action of genes ^118^. We tested genetic relationships only for cluster-pairs where both clusters exhibited significant SNP-heritability (p < .05; pre-corrected; 78 tests for area, 48 for thickness). Significance of genetic correlations was assessed at *p(FDR)* <.05.

#### 4.3.5 Associations with Cognition, Sex, Handedness, & ICV

Finally, we assessed relationships between asymmetry in our robust clusters and general cognitive ability, handedness, sex and ICV. For general cognition, we used the first principal component across the following 11 core UK Biobank cognitive variables ^95^: Mean reaction time (log transformed) [field 20023], Numeric memory [4282], Fluid reasoning [20016], Matrix completion [6373], Tower rearranging [21004], Symbol digit substitution [23324], Paired associate learning [20197], Prospective memory [20018] (recoded as 1 or 0, depending on whether the instruction was remembered on the first attempt or not), Pairs matching (log) [399], Trail making A (log) [6348], Trail making B (log) [6350]. Prior to the PCA, for participants with cognitive data, data was imputed for missing cognitive variables via the “imputePCA” R function (number of estimated components tentatively optimized using general cross validation; “missMDA” Package ^119^). PC1 (explaining 39.2%; Supplementary file 1L) was inversed to correlate negatively with age (r = -.39), ensuring higher values reflected higher cognition. As fewer participants had cognitive data relative to the other variables, for each cluster we ran one set of linear models to assess the marginal effect of cognition (PC1 as predictor; age, sex, ICV controlled; N = 35,199), and one set of linear models to assess the marginal effects of Handedness, Sex, and ICV in a model including all three predictors (age controlled, N = 37,570 with available handedness data). For the cognitive analysis, effects identified in the imputed dataset were checked against the confidence intervals for the effect in the subset of the data with no missing cognitive variables (N = 4696). Participants who self-reported as mixed handed were not included ^22^. Significance was considered at Bonferroni-corrected *α* = *p* < 7.3^−5^ (.01/136 [34 clusters × 4]).

## Data sharing/availability

All summary-level maps are available at neurovault.org/XXXX (upon acceptance). All code underlying the main analyses is available at https://github.com/jamesmroe/PopAsym and on the Open Science Framework (upon acceptance). All derived source data underlying all figures is also available here and in Supplementary files 2-3. All datasets used in this work are openly available, with the exception of LCBC, where participants, which include many children, have not consented to share their data publicly online. Other datasets used in this work are available without restrictions and are not subject to application approval (DLBS; https://fcon_1000.projects.nitrc.org/indi/retro/dlbs.html; CC BY-NC; SALD; http://fcon_1000.projects.nitrc.org/indi/retro/sald.html; CC BY-NC; IXI; https://brain-development.org/ixi-dataset; CC BY-SA 3.0). Accordingly, we have made the individual-level data for these samples available and our code can be used to reproduce vertex-wise analyses in these samples. Individual-level data for the remaining samples (LCBC; Cam-CAN, HCP; UKB) may be available upon reasonable request, given appropriate ethical, data protection, and data-sharing agreements where applicable. Requests must be submitted and approved via the relevant channel (details are provided in Supplementary File 1).

## Acknowledgements

Scripts were run on the Colossus processing cluster at the University of Oslo, and on resources provided by UNINETT Sigma2 (NN9769K). LCBC funding: European Research Council under grants 283634, 725025 (to A.M.F.), and 313440 (to K.B.W.); Norwegian Research Council (to A.M.F. and K.B.W.) under grants 249931 (TOPPFORSK) and 302854 (FRIPRO; to Y.W.), The National Association for Public Health’s dementia research program, Norway (to A.M.F). Data used in the preparation of this work were obtained from the MGH-USC Human Connectome Project (https://ida.loni.usc.edu/login.jsp). Data used in this work was also provided by the Cambridge Centre for Ageing and Neuroscience (CamCAN). This research has been conducted using the UK Biobank Resource.

