## Supplementary material for "Tracing the development and lifespan change of population-level structural asymmetry in the cerebral cortex": figure supplement

Roe et al.

Supplementary figures

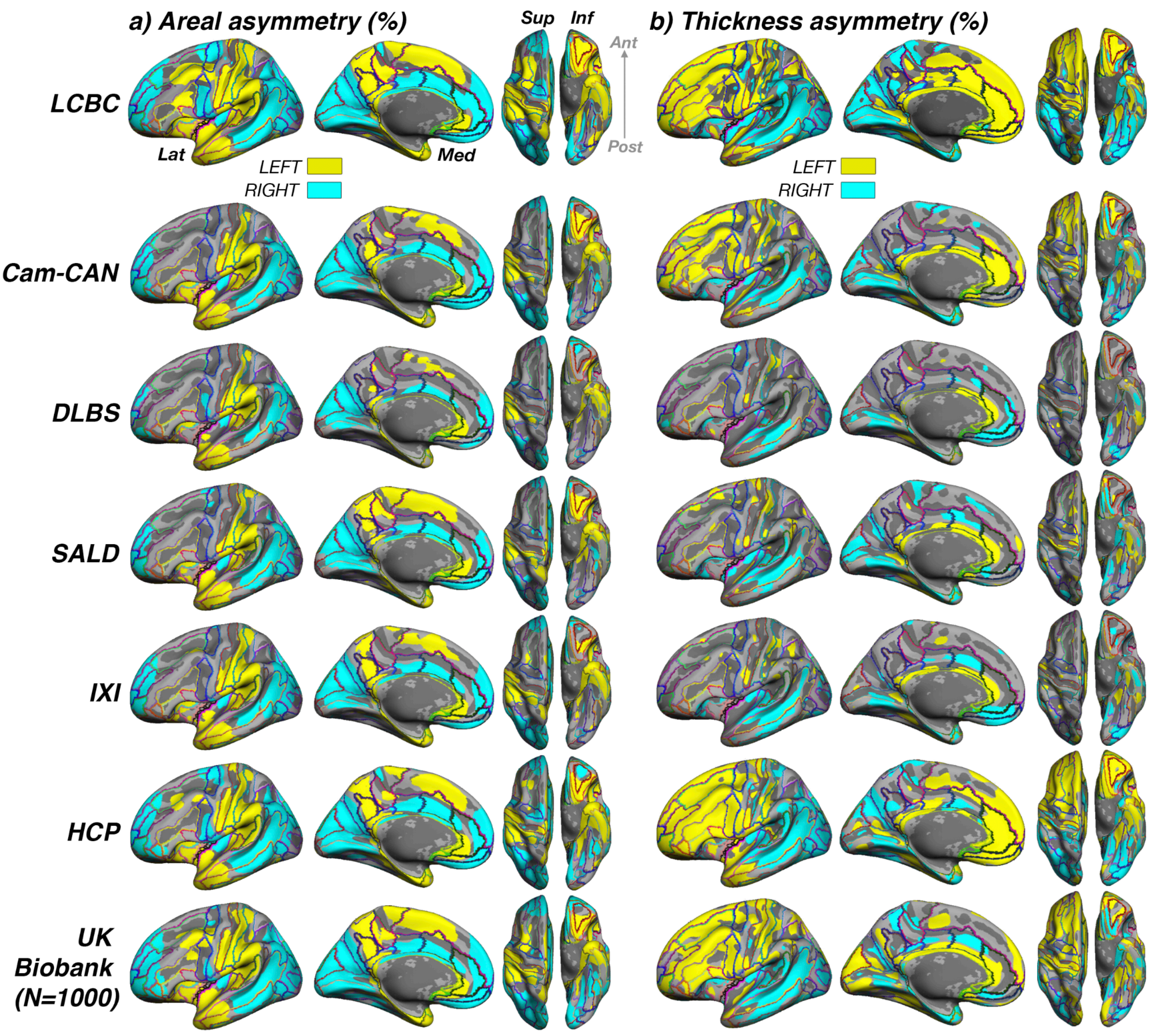


***Figure 1–figure supplement 1***

***Significance of asymmetry effects across samples***

*__A)__ Significance for __A)__ areal and __B)__ thickness asymmetry across all samples. Note that because differences in sample size and test power affect the overall level of significance and consequently the FDR-correction, the visualization threshold is set to match the FDR-corrected significance level for each cortical metric in the first sample (LCBC; p < .001). Warm and cold colours depict significance of leftward and rightward asymmetry, respectively. To permit more fine grained interpretation of anatomical correspondence, an outline of the Destrieux [1] cortical atlas is overlain. Compare with effect sizes in Fig. 1 and 2 in main paper.*

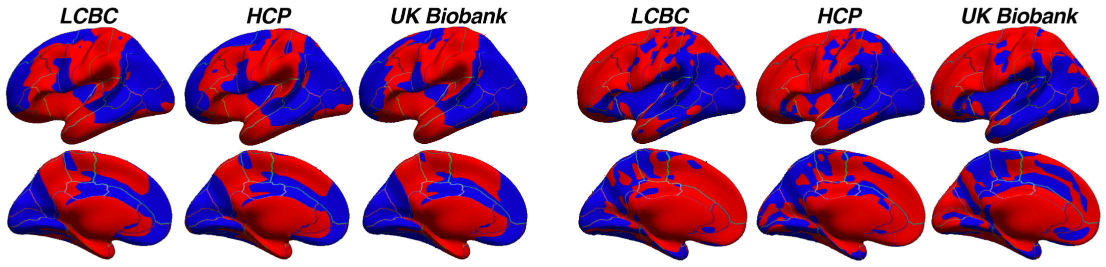


***Figure 1–figure supplement 2***

***Unthresholded maps***

*Completely unthresholded significance maps of areal (leftmost 3 columns) and thickness (rightmost 3 columns) asymmetry "effects" for the three largest samples. Desikan-Killiany atlas is overlain.*

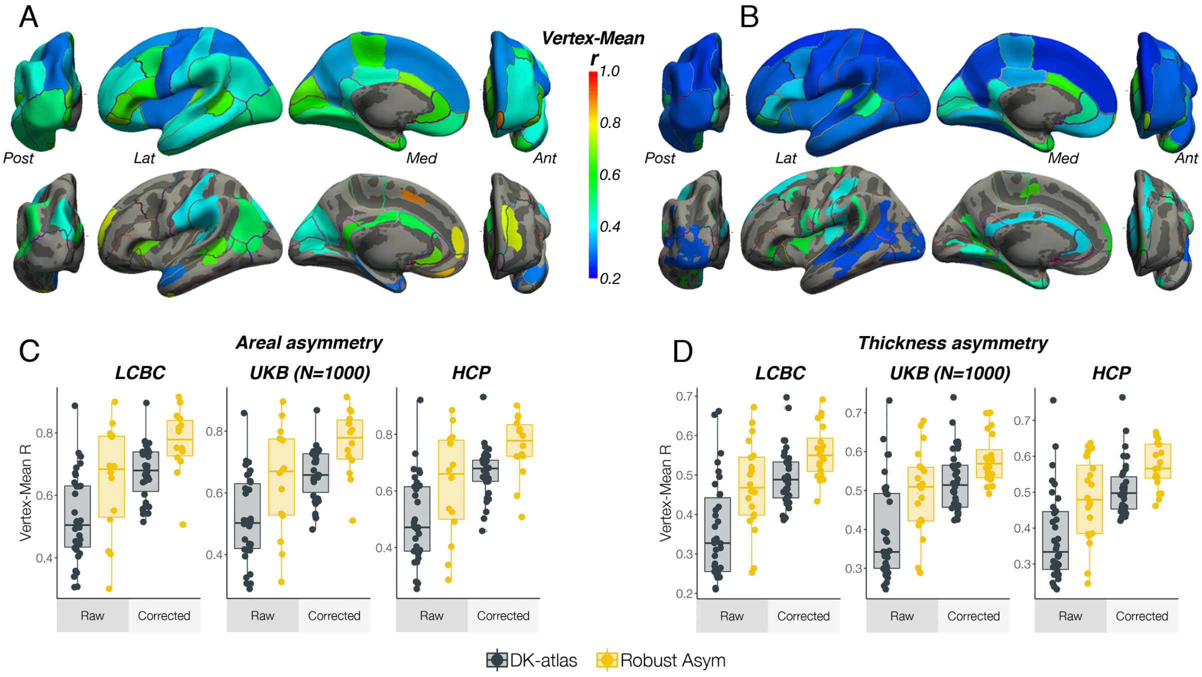


***Figure 1–figure supplement 3***

***Comparison of vertex-wise and atlas-based asymmetry estimates***

*To assess to what extent vertex-wise areal and thickness asymmetry estimates adhere to the anatomical boundaries of the Desikan-Killany (DK) atlas, we derived AI maps per subject and extracted a vertex × subject matrix. Then, within each of the 34 DK parcels we correlated the AI at each vertex with the parcel mean (i.e. the mean asymmetry across parcel vertices) and computed the mean correlation for each parcel. High correlations would be expected where atlas-derived parcels fit well to the underlying vertex-wise structure of cortical asymmetry. We then repeated this analysis using our set of robust asymmetry clusters. As a formal test, for each cortical metric the resulting coefficients were used as response variable in linear regressions with Cluster Type (DK parcels vs. Robust clusters) as predictor, controlling for cluster size (nVertices) and the Cluster Type × nVertices interaction. For areal asymmetry, the average vertex-mean correlations across all Desikan-Killiany (DK) parcels were $r = .53 \pm .14$ [LCBC], $r = .51 \pm .14$ [UK Biobank], and $r = .49 \pm .15$, which respectively increased to $r = .65 \pm .18$, $r = .64 \pm .18$, and $r = .62 \pm .19$ for areal asymmetry clusters. For thickness, the average vertex-mean correlations across all DK parcels were $r = .36 \pm .12$ [LCBC], $r = .39 \pm .12$ [UK Biobank] and $r = .38 \pm .12$, which respectively increased to $r = .47 \pm .11$, $r = .50 \pm .11$ and $r = .48 \pm .12$. As would be expected, within parcel/cluster vertex-mean correlations were significantly lower in larger parcels/clusters. Linear regressions (size controlled) revealed a significant main effect of Cluster Type in all but one association test (see Supplementary file 1D), confirming the visual impression that DK parcels conform poorly to the underlying asymmetry of cortex.* ***A-B)*** *Average correlation between vertex-wise estimates asymmetry within DK parcels to the mean across each parcel (top rows) and between vertex-wise asymmetry estimates within robust clusters to the mean across each cluster (bottom rows), for areal (A) and thickness (B) asymmetry. The results on the surface are the average across the LCBC, UKB, and HCP datasets used in the vertex-wise* *analysis in main paper (the complete set of raw vertex-mean correlation coefficients correlated highly between all dataset-pairs for both areal [min r = 0.97] and thickness asymmetry [min r = 0.95]***. C-D)** *Vertex-mean correlation coefficients for areal asymmetry in DK parcels and robust asymmetry clusters, shown as raw values and after correcting for number of vertices in the parcel/cluster, for areal (C) and thickness asymmetry (D). Lat=lateral; Med=medial; Post=posterior; Ant=anterior.*

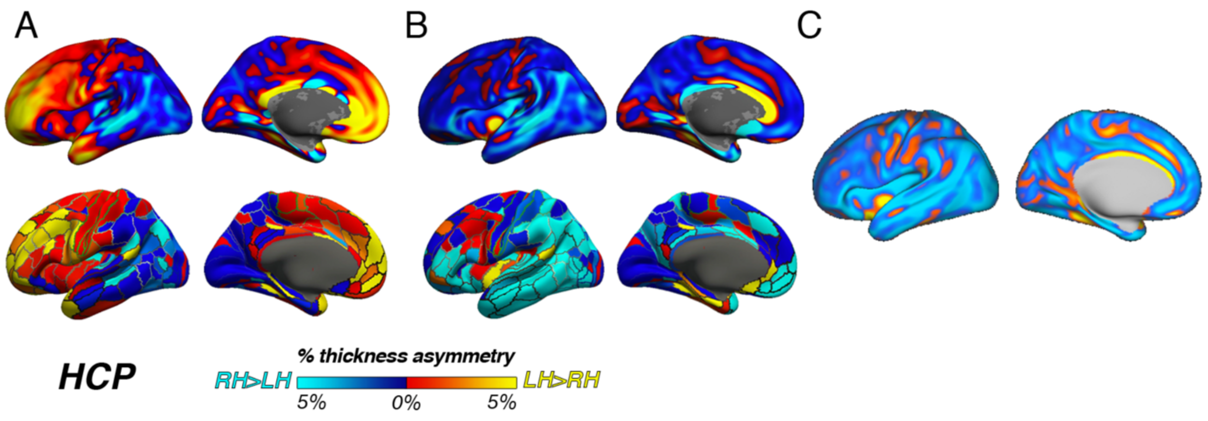


***Figure 1–figure supplement 4***

***HCP pipeline***

*Thickness asymmetry results in the HCP dataset vary depending on the preprocessing pipeline. A) Results from using the –hires argument to recon-all (as in main paper). We used this method to best harmonize preprocessing across cohorts whilst accounting for the higher resolution of HCP data. Shown again for comparison, the top row in A is akin to the results in the main paper (analysed using cross-hemispheric registration methods) whereas the bottom row is the same data analysed using standard parcellation methods (as in Figure 1–figure supplement 5). B) Unthresholded thickness asymmetry results using the HCP preprocessed data subject to extra preprocessing steps and inputs, analysed using cross-hemispheric registration methods (top row) and standard parcellation methods (bottom). C) Results using the HCP preprocessed data when calculating thickness asymmetry on the fs_LR template. Warm and cold colours depict leftward and rightward asymmetry, respectively.*

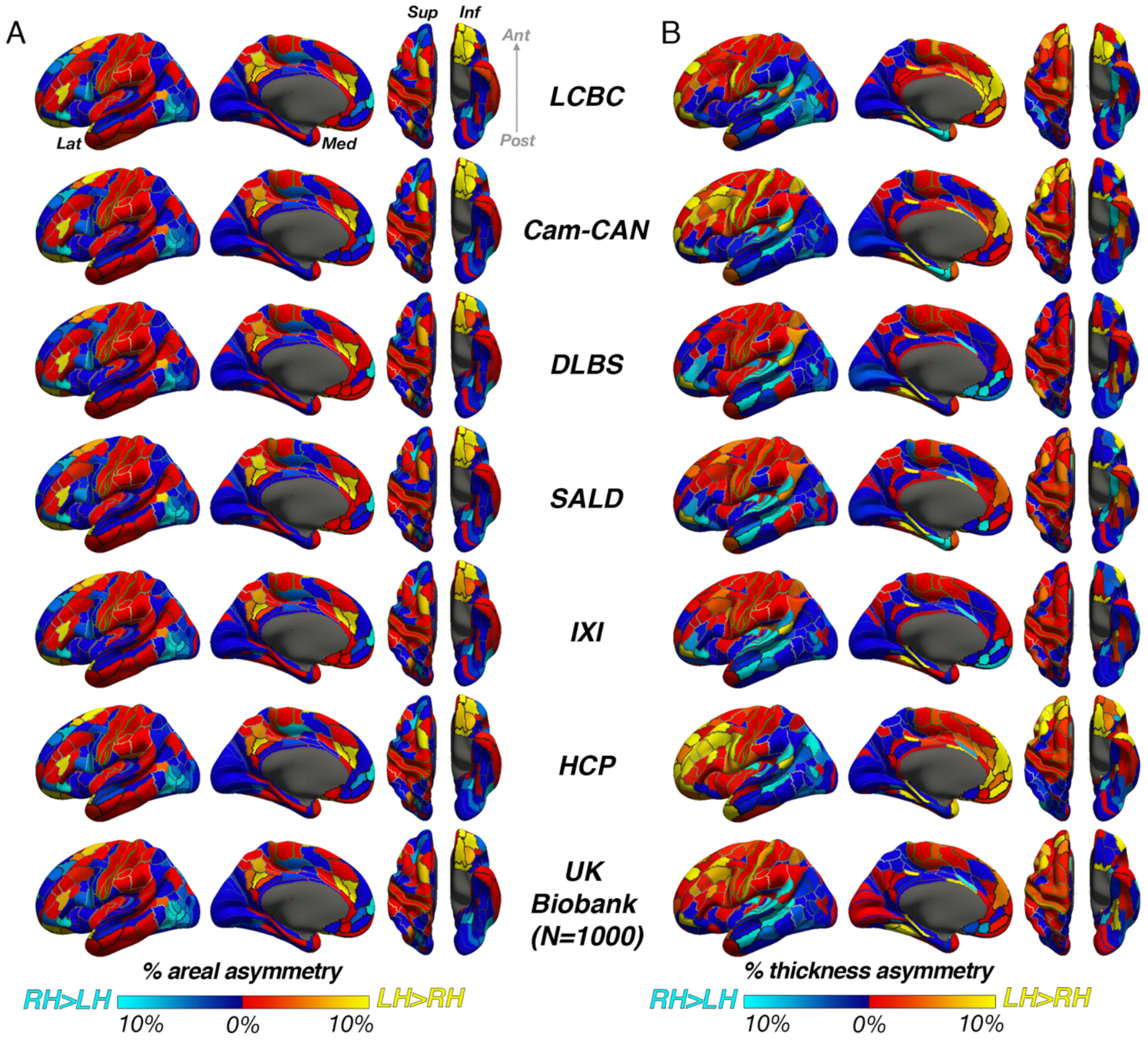


***Figure 1–figure supplement 5***

***Unthresholded asymmetry effects analysed using a standard brain atlas with no cross-hemispheric registration of data***

*Mean areal and B) thickness asymmetry in each dataset analysed using standard methods on the fsaverage template within parcels from the HCP multimodal brain atlas [2]. We used this atlas here because it appeared best suited to assess parcels that are homotopic. Warm and cold colours depict leftward and rightward asymmetry, respectively. Post=posterior; Lat=lateral; Med=medial; Ant=anterior; Sup=superior; Inf=inferior.*

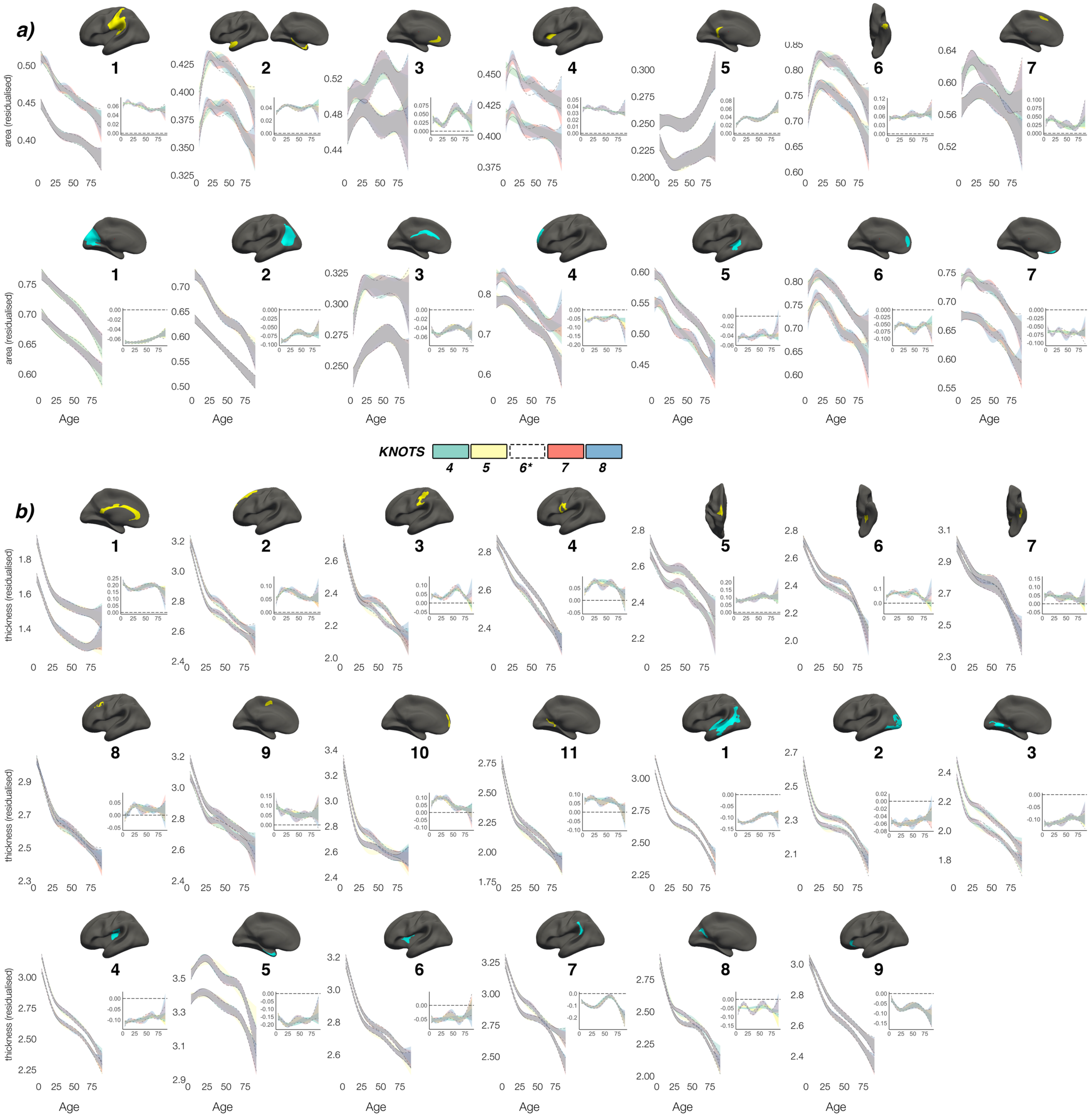


***Figure 3–figure supplement 1***

***Knot comparison***

*Lifespan trajectories of population-level* ***a)*** *areal and* ***b)*** *thickness asymmetries modelled with different smoothing parameters (number of knots 4-8, with the selected number of knots [6] shown in black dotted outline). Grey indicates* *overlap. All age trajectories were fitted using GAMMs. Inset plots show absolute asymmetry trajectories across life at the different smoothing parameters.*

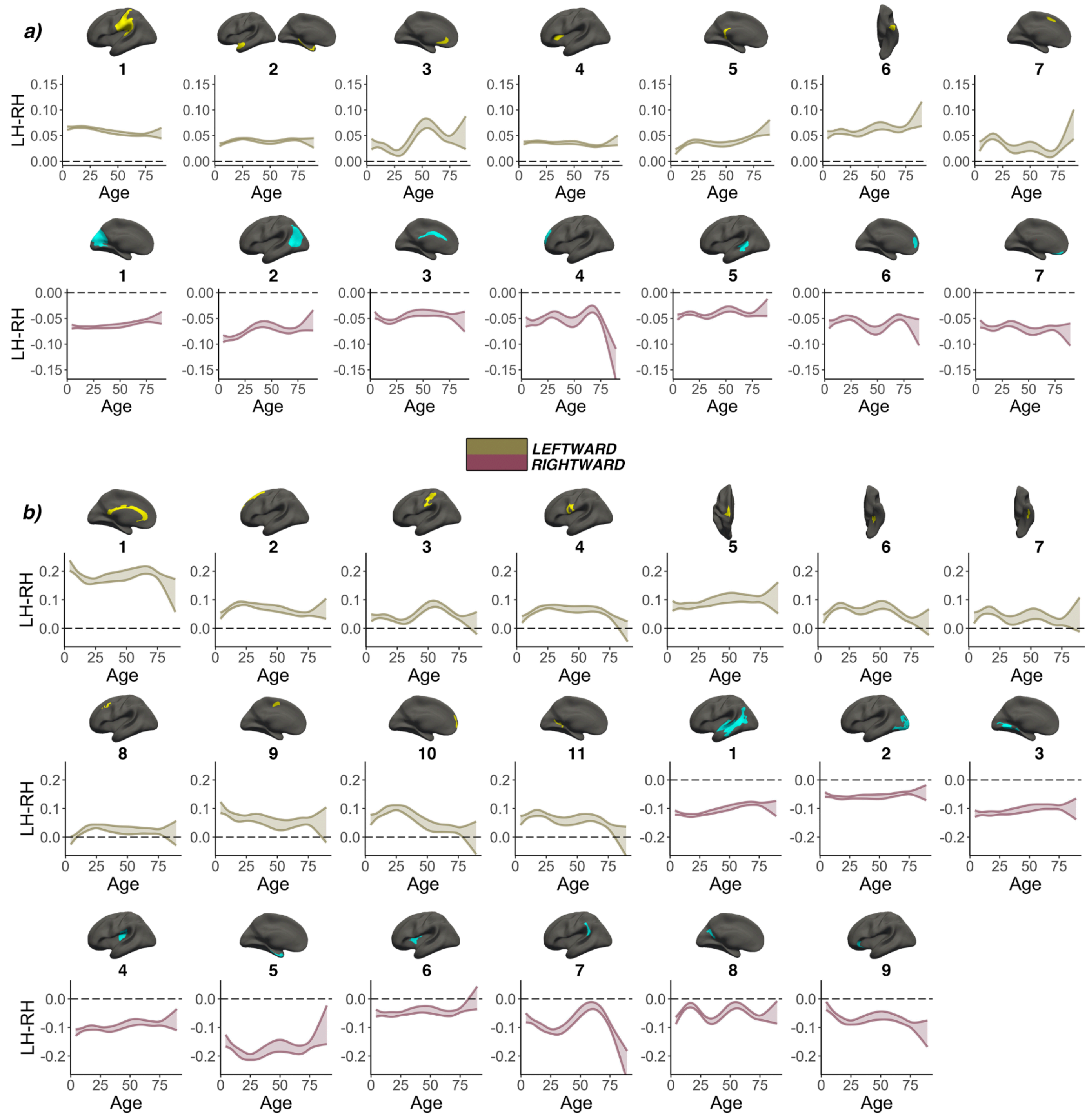


***Figure 3–figure supplement 2***

***Smooth Age x Hemisphere interactions***

*Generalized additive mixed Model (GAMM) results for population-level **a)** areal and **b)** thickness asymmetries. A smooth Age × Hemisphere interaction [s(LH)-s(RH-Age)] was fitted to model asymmetry changes across the lifespan. Specifically, GAMMs were used to compute the zero-centered age-trajectories of the left [s(LH-Age)] and right [s(RH-Age)] hemisphere for each cluster, and the asymmetry trajectory was computed as the difference between the two [s(LH-Age)-s(RH-Age)]. Gold and pink colours denote clusters defined by leftward or rightward asymmetry, respectively. For each cluster the main effect of Hemisphere was added to visualize the lifespan trajectory of absolute asymmetry. Bands represent 95%*

*confidence intervals. Note that all areal clusters show asymmetry trajectories that are significantly different from 0 (symmetry; dotted line) across the entire lifespan, and 19/20 thickness clusters were significantly asymmetric already by ~age 4.*

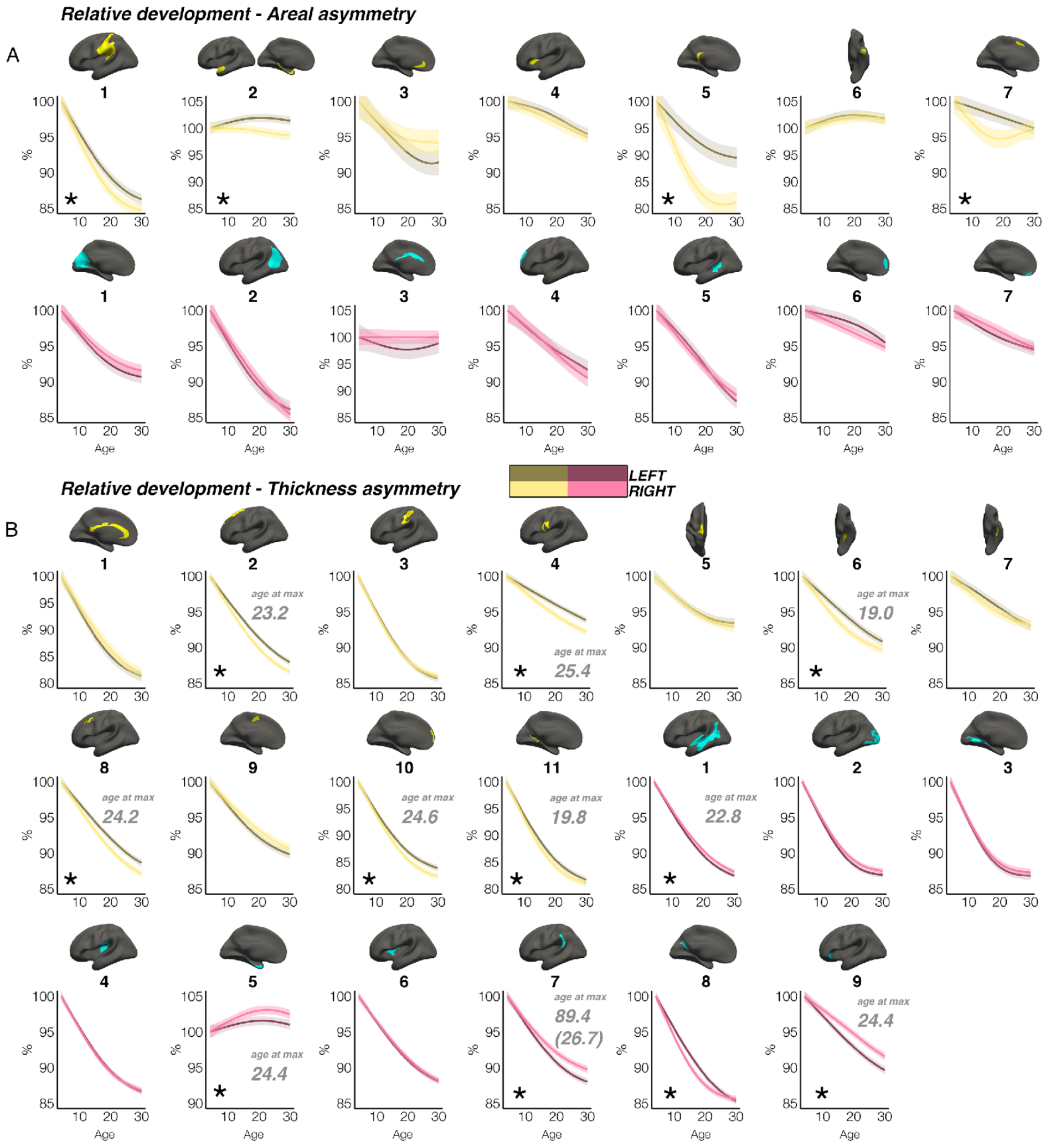


***Figure 3–figure supplement 3***

***Relative developmental trajectories***

*Relative developmental trajectories of population-level areal (A) and thickness asymmetries. (B). Relative change was calculated by scaling the LH and RH fitted lifespan trajectories by the prediction at the minimum age (i.e. ~4 years). To highlight development, the x-axis covers the age-range 4-30 years, although the results are calculated* *from full lifespan models. Darker trajectories indicate LH trajectories. Shaded areas indicate 95% CI. Relative developmental change between hemispheres is suggested if the CI's of relative LH and RH trajectories diverge to be non-overlapping (denoted with *). Note that for some areal asymmetries with suggestive relative hemispheric development, the data nevertheless was more indicative of stable and parallel trajectories (e.g. L1 and L2; see Figures 2 & 3), suggesting these indications of relative developmental hemispheric differences should be interpreted cautiously and in combination with the full lifespan GAMM trajectories shown in Figures 2 & 3, which contain inherent noise possibly due to the complexity of lifespan data (see Limitations). Note also that we report 10/20 relative developmental hemispheric differences for thickness asymmetry, as cluster R8 appeared to exhibit an initial faster thinning of the right hemisphere despite maintaining rightward thickness asymmetry throughout development (compare with Figure 3B). Where a relative developmental hemispheric difference was evident for thickness asymmetry, age at the point of maximum relative asymmetry across life is given, denoted in grey (calculated as the age of maximally non-overlapping CI's). One cluster (R7) appeared to exhibit maximum thickness asymmetry in the estimated trajectories at age 89.4 (see also Figure 3), so we additionally calculated age at maximum asymmetry for the first developmental peak (in parentheses). All lifespan age trajectories here were fitted using equivalent GAMMs as in the main paper, with the addition of a (scaled) ICV covariate (i.e.* `gamm(Y ~ s(Age, by = as.factor(hemi), k = 6) + as.factor(hemi) + Sex + Scanner + ICV, data = DF, random = ~ (1 | ID))`).

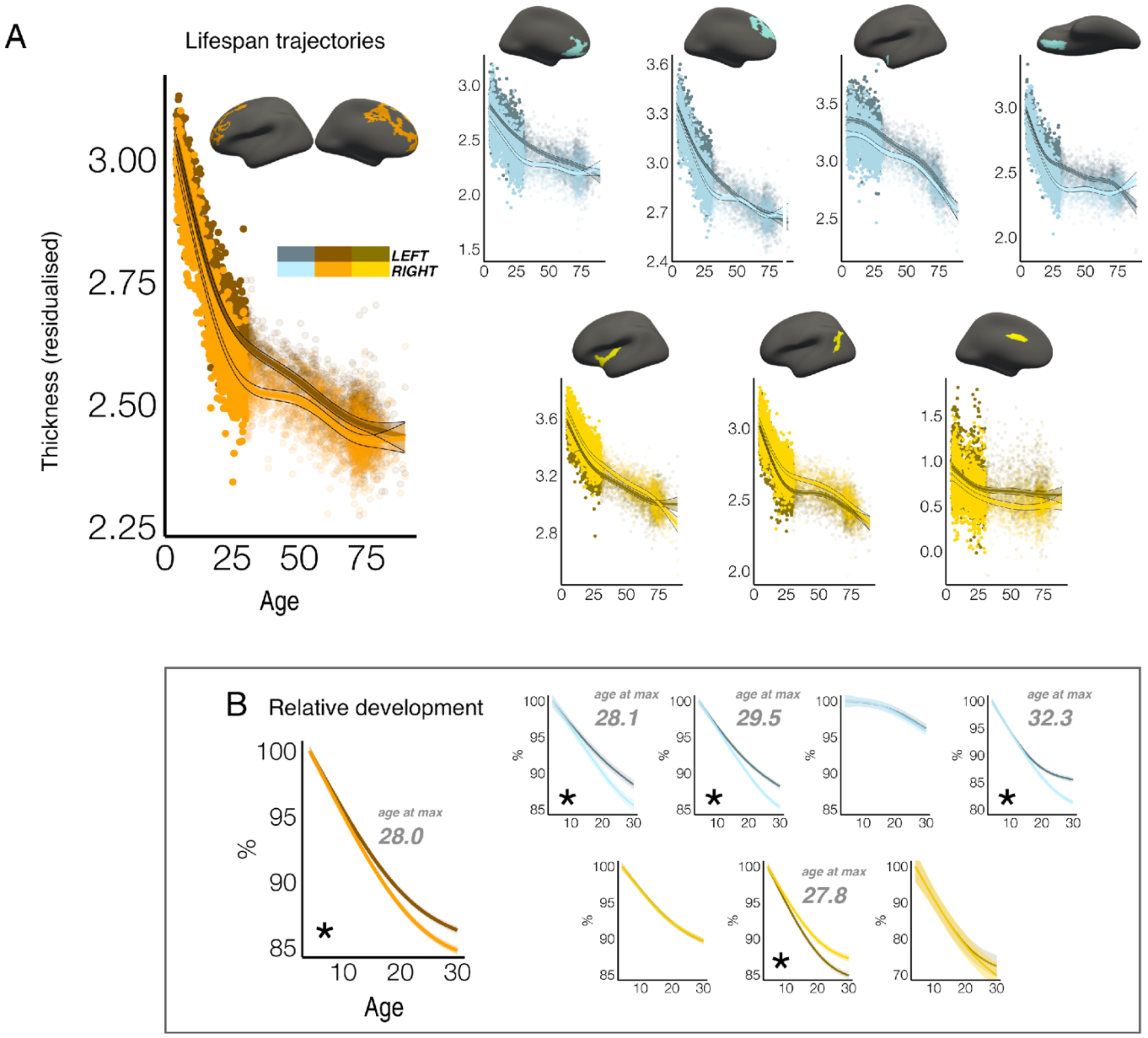


***Figure 3–figure supplement 4***

***Lifespan thickness trajectories in regions exhibiting age-related change in asymmetry***

***A)*** *Homotopic lifespan thickness trajectories in an alternative set of regions derived from a previous analysis (clusters from Roe et al., 2021 [3]; derived from vertex-wise analyses of age-related thickness asymmetry change in LCBC adult data [20-89 years]). Dark colours correspond to LH trajectories (colours correspond to clustering solutions in Roe et al., 2021 [3]). To highlight development, datapoints are semitransparent after age 30. Shaded areas indicate 95% CI.* ***B)*** *Relative developmental LH and RH thinning from age 4-30. Plots correspond to the brain regions shown in A. Relative change was* *calculated by scaling the LH and RH fitted trajectories by the prediction at the minimum age (i.e. ~4 years). To highlight development, the x-axis covers the age-range 4-30 years, although the results are calculated from full lifespan models. Relative developmental change between hemispheres is suggested if the CI's of relative LH and RH trajectories diverge to be non-overlapping (denoted with *). These indications of relative developmental hemispheric differences should be interpreted cautiously and in combination with the full lifespan GAMM trajectories shown in A, which contain inherent noise possibly due to the complexity of lifespan data (see Limitations). Where a relative developmental hemispheric difference was evident for thickness asymmetry, age at the point of maximum relative asymmetry across life is given, denoted in grey (calculated as the age of maximally non-overlapping CI's). All lifespan age trajectories here were fitted using equivalent GAMMs as in the main paper, with the addition of a (scaled) ICV covariate* (i.e. `gamm(Y ~ s(Age, by = as.factor(hemi), k = 6) + as.factor(hemi) + Sex + Scanner + ICV, data = DF, random = ~ (1 | ID))`).

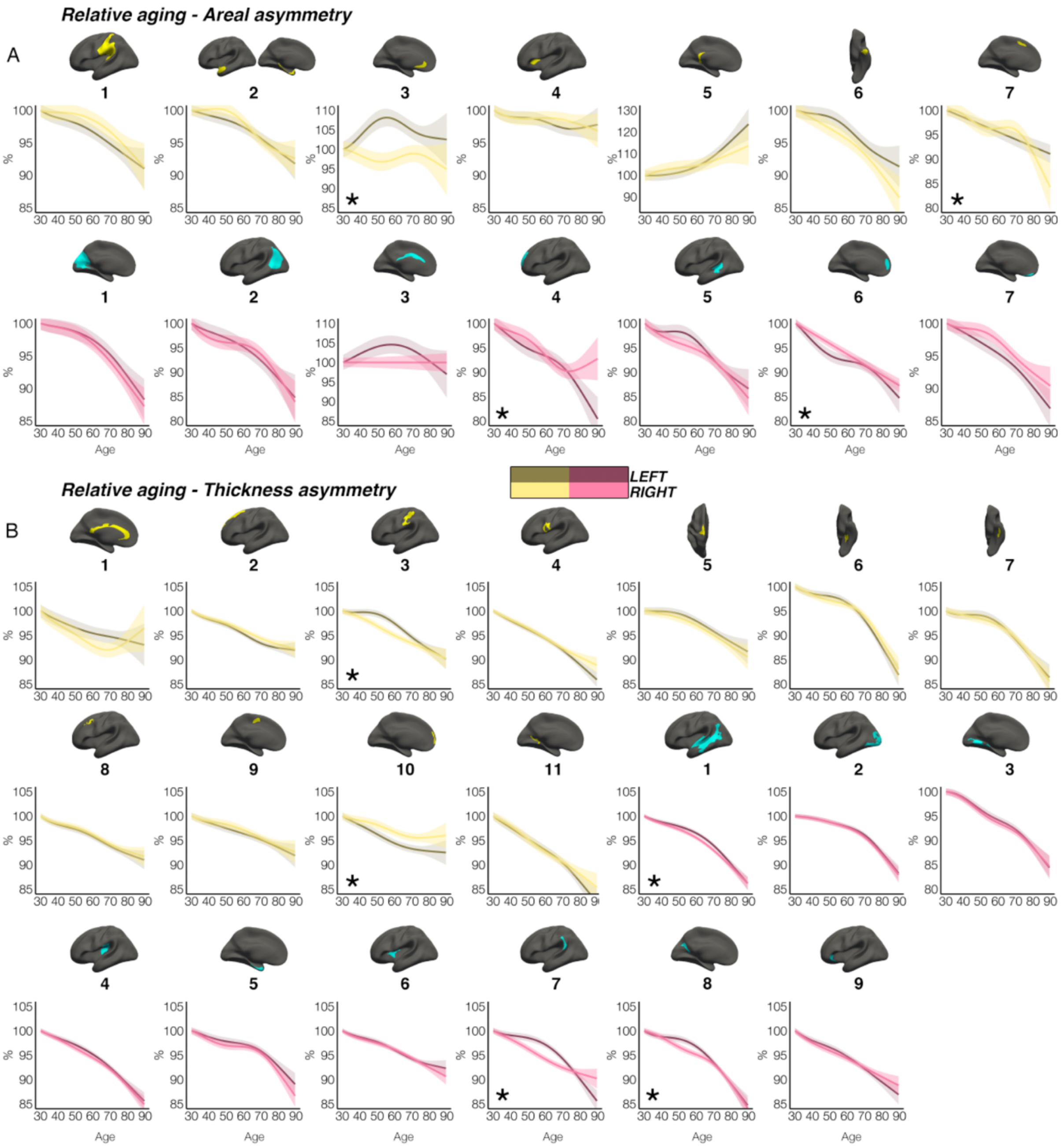


***Figure 3–figure supplement 5***

***Relative aging trajectories***

*Relative aging trajectories of population-level areal (A) and thickness asymmetries. (B). Relative change was calculated by scaling the LH and RH fitted lifespan trajectories by the prediction at age 30 years. To highlight aging, the x-axis covers the age-range 30-89 years, although the results are calculated from full lifespan models. Darker trajectories indicate LH trajectories. Shaded areas indicate 95% CI. Relative aging-related change between hemispheres is suggested if the CI's of relative LH and RH trajectories diverge to be non-overlapping (denoted with *). These indications of relative aging-related hemispheric differences should be interpreted cautiously and in combination with the full lifespan GAMM trajectories shown in Figures 2 & 3. All lifespan age trajectories here were fitted using equivalent GAMMs as in the main paper, with the addition of a (scaled) ICV covariate* (i.e. `gamm(Y ~ s(Age, by = as.factor(hemi), k = 6) + as.factor(hemi) + Sex + Scanner + ICV, data = DF, random = ~ (1 | ID))`).

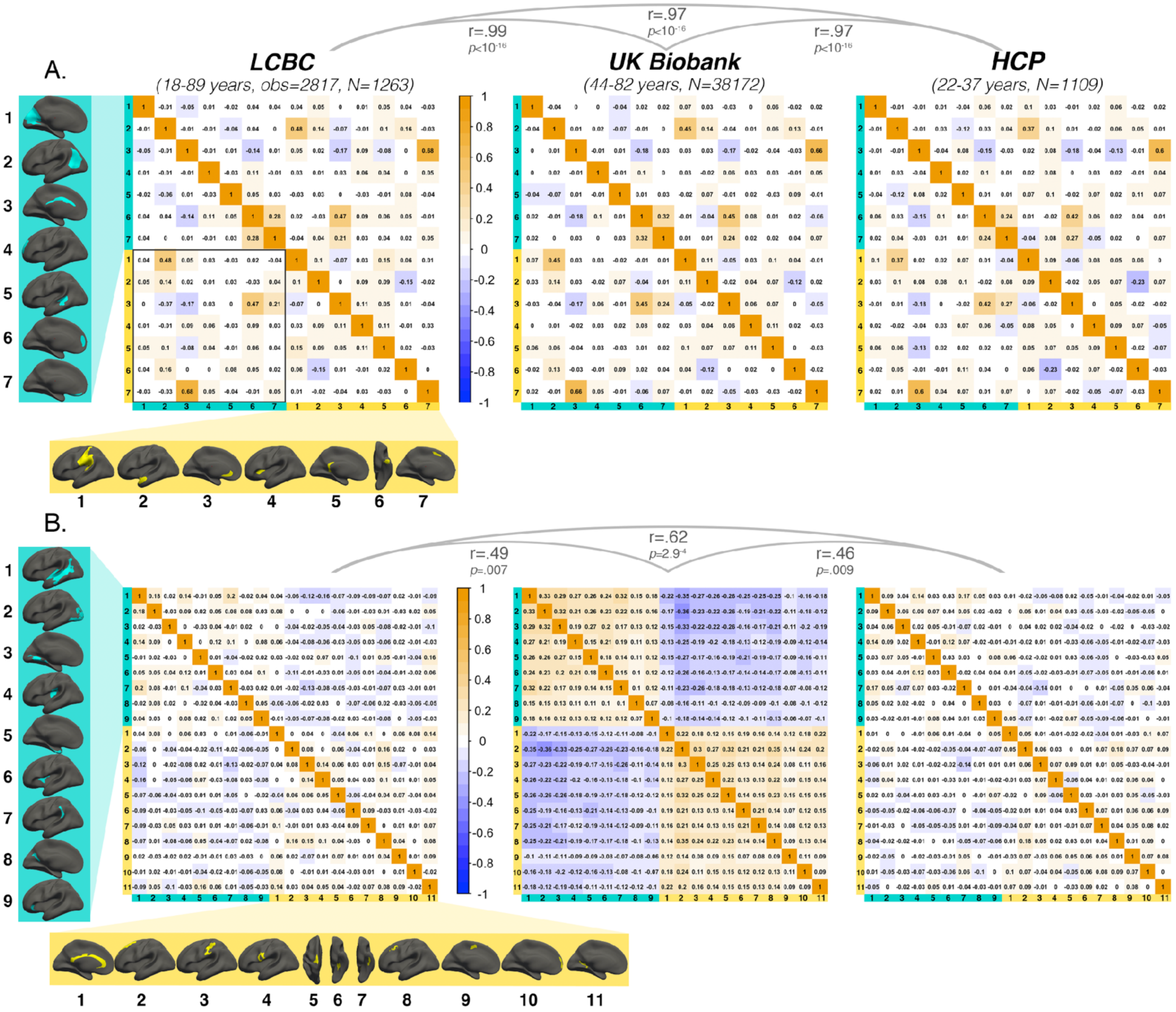


**Figure 5–figure supplement 1**

**Annotated covariance matrices**

*Figure 4: Interregional correlations between A) areal asymmetries and B) thickness asymmetries for each replication dataset (AI's residualised for age, sex, scanner). AI's in rightward clusters are inversed, such that positive correlations denote positive asymmetry-asymmetry relationships, regardless of direction of mean asymmetry in the cluster (i.e. higher asymmetry in the population-direction). Yellow and blue brain clusters/colours denote leftward and rightward asymmetries, respectively (clusters numbered for reference). A consistent covariance structure was evident both for areal (r >= .97) and thickness asymmetry (r >=.49; results above matrices). Black box in A highlights relationships between opposite-direction asymmetries (i.e. leftward vs rightward regions). Interregional correlations for thickness asymmetry in the lower left quadrant of UK Biobank matrix are visualized in Figure 5–figure supplement 2.*

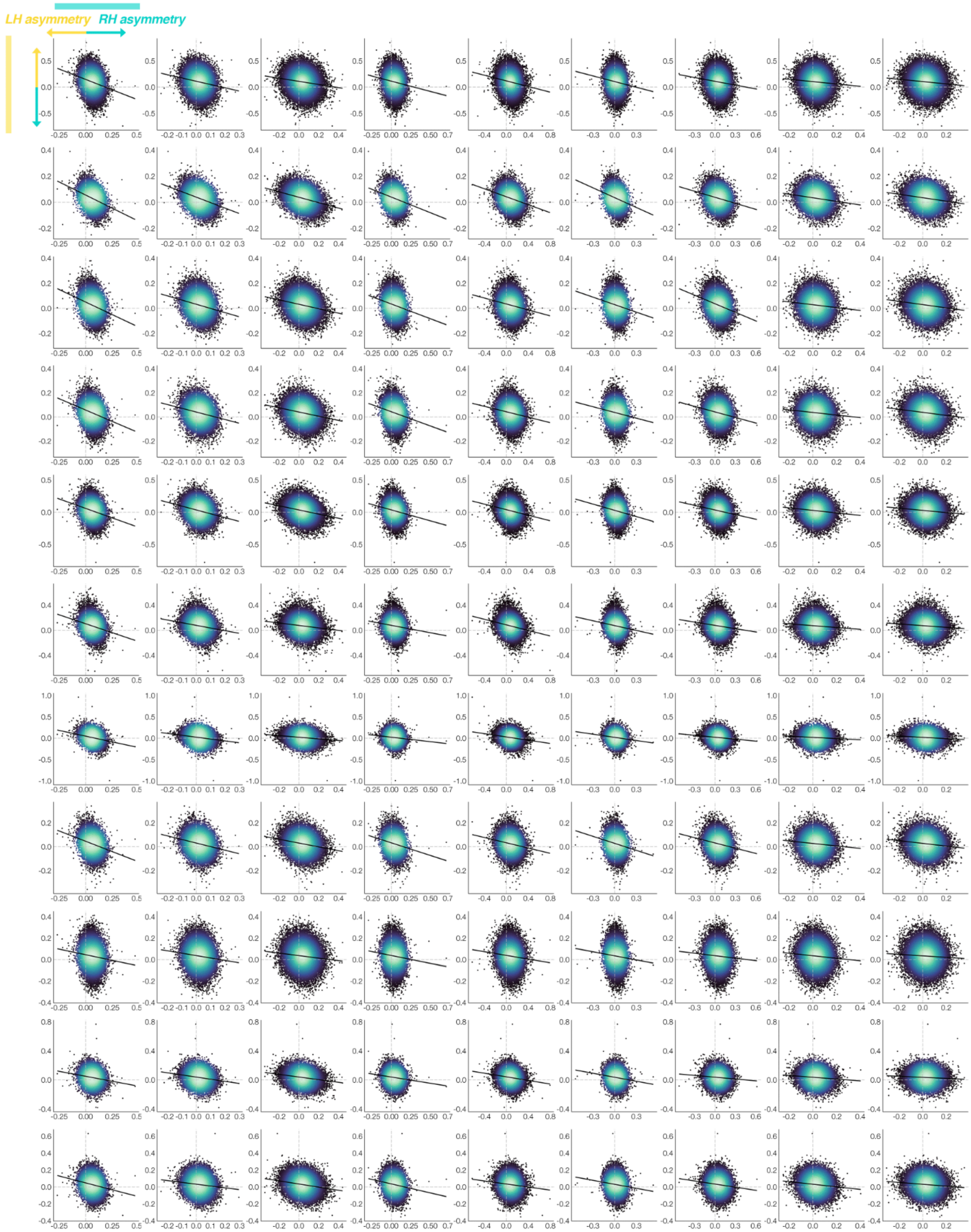

LH asymmetry
RH asymmetry

***Figure 5–figure supplement 2***

***SI Fig. 10 – UK Biobank lower quadrant***

*Thickness asymmetry interregional correlations between opposite-direction asymmetries (lower left quadrant of the UK Biobank correlation matrix in SI. Fig. 3; N = 38,172) are visualized to describe whether negative asymmetry-asymmetry correlations pertained to reduced or reversed thickness asymmetry. Lines of symmetry (0) are shown in grey. X-axis and Y-show raw AI data after removing the fixed effects of age and sex. AI's in rightward clusters are inversed, such that positive correlations would denote positive asymmetry-asymmetry relationships regardless of direction of mean asymmetry in the cluster (i.e. higher asymmetry in the population direction). Order of cortical locations is in Figure 5–figure supplement 1.*

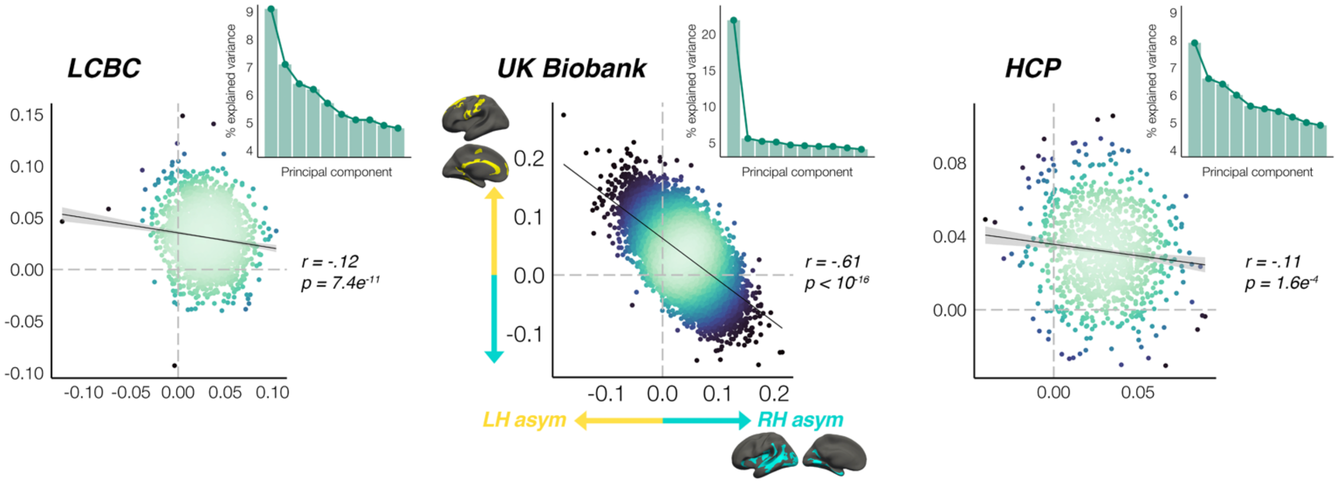


***Figure 5–figure supplement 3***

***SI Fig. 11 – Global thickness asymmetry relationships***

*Global thickness asymmetry relationships. Principal components analysis across AI's in all leftward and rightward thickness asymmetry clusters revealed a single component explained most of the variance in UK Biobank, and there was evidence of a relatively stronger first component in LCBC and HCP (inset scree plots). Scatter plots show the partial correlation of the mean asymmetry across all leftward vs. all rightward clusters (means weighted by cluster size) plotted for each cohort, after AI's were corrected for age, sex and (where applicable) scanner. Lines of symmetry (0) are shown as dotted grey. AI's in rightward clusters are inversed, such that positive correlations would denote positive asymmetry-asymmetry relationships regardless of direction (i.e. higher asymmetry in the population direction).*

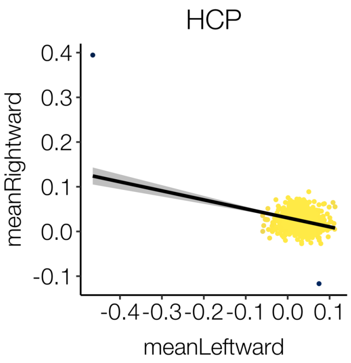


***Figure 5–figure supplement 4***

***HCP outliers discarded***

*Two outliers (black) were detected in the cortical thickness data of HCP and subsequently discarded for all analyses following the vertex-wise delineation of cortical asymmetry (in which their inclusion had negligible effect on the derived mean asymmetry maps). Plot shows mean thickness asymmetry across all leftward vs. mean thickness asymmetry across all rightward clusters.*

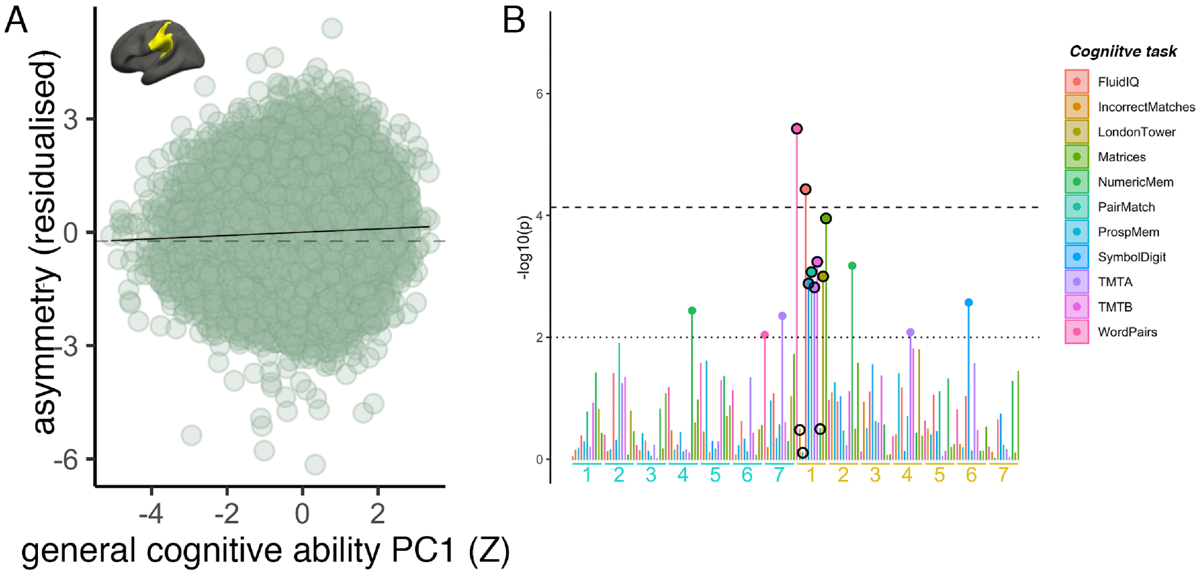


***Figure 7–figure supplement 1***

***A)*** *Association of reduced leftward areal asymmetry in the large supramarginal cluster upon general cognitive ability (PC1 across 11 cognitive tests). The line of null association is shown for comparison (dotted).* ***B)*** *Post-hoc association tests between cognitive scores on the 11 separate tests in UKB and areal asymmetry in our 14 robust clusters (labelled under plot, leftward and rightward clusters in yellow and blue text, respectively). Black circled points highlight the association tests for the leftward cluster (visualized) wherein we find the association with general cognitive ability (i.e. PC1). Corrected [p < 7.3e$^{-5}$] and uncorrected threshold [p = .01] shown by dotted and non-dotted line, respectively. As these tests are a post-hoc confirmation and not independent of the initial analysis PC1 across tasks, the correction level is the same as in the main paper (though we note the association with PC1 survives regardless [p = 7.33$^{-7}$]).*

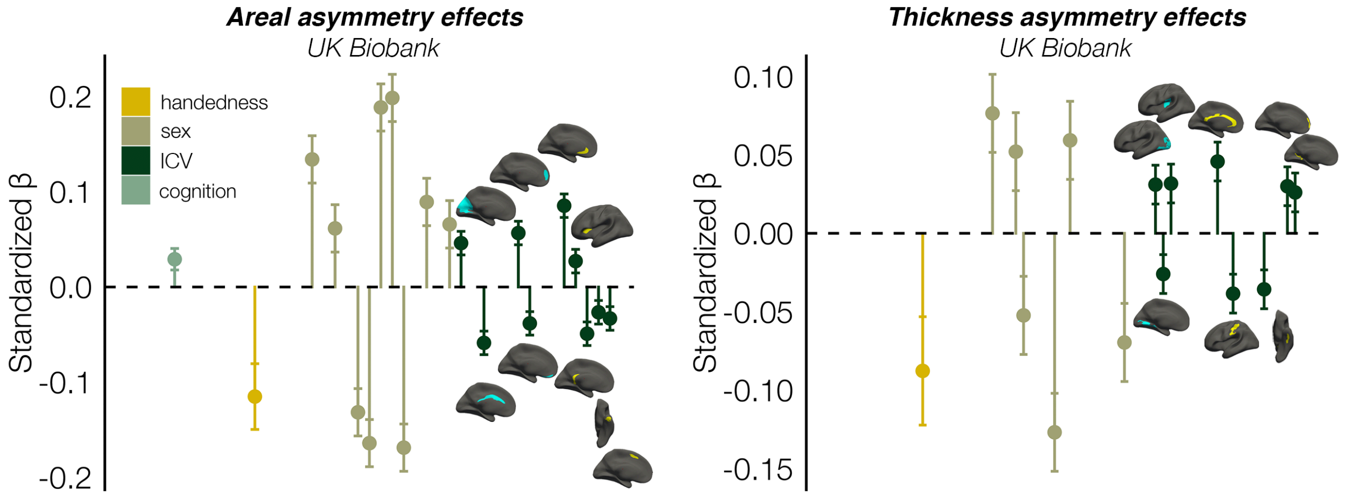


**Figure 7–figure supplement 2**

**ICV effects**

*Continuation of Fig. 5 in main paper. Cortical locations of the effects underlying the observed associations of robust areal (left) and thickness (right) asymmetry phenotypes with brain size (ICV) in UK Biobank. Plots denote the effect sizes (standardized Betas) and cortical location of all associations surpassing Bonferroni-corrected significance (Fig. 5). Right handers and females are coded 0, such that a negative effect size for handedness / sex / ICV / cognition denotes less asymmetry in left handers / males / larger brains / higher cognition. Blue and yellow clusters denote leftward and rightward asymmetry, respectively.*
